# Whole-Genome Sequencing, Annotation and Phenotypic Taxonomic Confirmation of a Multidrug-Resistant *Escherichia coli* Strain Isolated from the Blood of a Sepsis Patient

**DOI:** 10.1101/2025.03.21.644136

**Authors:** Peter M. Eze, Lucy Dillon, Jurnorain Gani, Timothy Planche, Linda B. Oyama, Christopher J. Creevey

**Affiliations:** School of Biological Sciences, Queen’s University Belfast, United Kingdom; Department of Environmental Health Science, Nnamdi Azikiwe University, Nigeria; Institute of Infection and Immunity, City St. George’s, University of London, United Kingdom

**Keywords:** *Escherichia coli*, sepsis, antimicrobial resistance, whole-genome sequence, phenotypic, genomic

## Abstract

Sepsis (blood stream infection) caused by multidrug-resistant (MDR) bacteria, particularly *Escherichia coli*, represents a significant global health threat due to high morbidity, mortality, and limited treatment options. *E. coli*, a major causative agent of bloodstream infections, has evolved highly virulent and MDR strains, which contribute to the increasing burden of antimicrobial resistance (AMR), complicating clinical management and reducing the efficacy of conventional antibiotic therapies. In this study, we characterised the genomic and phenotypic drug resistance mechanism of *E. coli* 266631E isolated from a sepsis patient, highlighting the negative implications of MDR *E. coli* in sepsis. Antimicrobial susceptibility testing and minimum inhibitory concentration analysis revealed resistance to multiple antibiotics, including amoxicillin, cefotaxime, ciprofloxacin, gentamicin, and tobramycin. Whole genome sequencing identified a broad array of AMR genes encoding resistance to various antibiotic classes, such as macrolides, fluoroquinolones, aminoglycosides, carbapenems, and cephalosporins. Notably, the CTX-M-15 gene, a key extended-spectrum β-lactamase determinant, was found in both the bacterial chromosome and an IncF-type plasmid, emphasizing the potential for horizontal gene transfer and rapid dissemination of resistance. Confirming the taxonomy of the novel and unidentified bacterial strain through querying its 16S rRNA sequence and genome in recognised bacterial taxonomic databases presented a challenge. The isolate showed genetic similarity to *E. coli, E. fergusonii*, and *Shigella* species despite their phenotypic differences and variations in their pathogenic traits. However, a simple phenotypic laboratory procedure, based on the biochemical and cultural differences among these bacteria in Coliform *ChromoSelect* Agar, confirmed the isolate as *E. coli*. This study underscores the critical importance of integrating phenotypic methods with genomic tools for the accurate identification of clinically significant bacteria. It also highlights the need for both phenotypic and genetic surveillance of key MDR variants in healthcare settings to enable timely, precise diagnosis and targeted treatment of life-threatening infections such as sepsis.

**DATA SUMMARY:** The outputs of the MALDI-TOF analysis, AMR analysis using AMRFinderPlus, starAMR, and RGI, the query of bacterial 16S rRNA in the NCBI, Greengenes2, and SILVA databases, as well as Sourmash and pangenome analyses, are available in the supplementary material. The bacterial 16S rRNA gene sequence has been deposited in the NCBI GenBank database under accession number PQ871642. The isolate’s complete genome sequence has been submitted to the NCBI Genome database under BioProject accession PRJNA1220687 and BioSample accession SAMN46722626. The chromosome is available under GenBank accession number CP183489, while plasmids and other contigs are available under accession numbers CP183490–CP183498. The scripts used in the following bioinformatics analysis – Sourmash, Roary, and Prokka (for pangenome analysis) are available at: https://github.com/LucyDillon/MDR_isolate.

**IMPACT STATEMENT:** This study presents a comprehensive genomic and phenotypic characterization of a multidrug-resistant *Escherichia coli* strain implicated in sepsis, uncovering extensive antimicrobial resistance genes across both the chromosome and plasmids. It exposes taxonomic ambiguity with closely related species such as *E. coli, Shigella*, and *E. fergusonii*, underscoring gaps in current genomic databases and reinforcing the necessity of phenotypic testing. By integrating whole-genome sequencing with biochemical differentiation, the research strengthens diagnostic precision and informs clinical decision-making for life-threatening infections. Overall, it advances AMR and bacterial taxonomy research by demonstrating the value of an integrated genomic-phenotypic framework to accurately identify and guide the treatment and surveillance of emerging MDR pathogens in clinical settings.

## INTRODUCTION

*E. coli* is an opportunistic pathogen but is typically a commensal within the gut microbiota of humans and other animals. It is one of the most versatile bacteria, capable of causing a variety of infections, including both intestinal and extraintestinal infections (Denamur et al., 2021; Priyanka et al., 2023). Extraintestinal pathogenic *E. coli* (ExPEC) comprises a group of strains with the ability to colonize and infect extraintestinal sites. Systemic infections caused by ExPEC are referred to as invasive *E. coli* disease (IED) or invasive ExPEC disease. This condition includes infections of the bloodstream and other normally sterile body sites, such as cerebrospinal fluid, the pleural cavity, peritoneal space, bone, and joints. Invasive *E. coli* disease may result in sepsis, septic shock, or death (Doua et al., 2023; Geurtsen et al., 2022; Rosenberg et al., 2021; Dugar et al., 2020). Colonization and infection of extraintestinal sites by pathogenic *E. coli* are leading causes of sepsis in both hospitals and communities (Markwart et al., 2020; Shao et al., 2023).

*Escherichia coli* is the most prevalent bacterium causing sepsis and one of the most common causes of bloodstream infections, as it is capable of accessing and surviving in the bloodstream (AL-Khikani et al., 2023; Shao et al., 2023). *E. coli* has been reported as the leading cause of bloodstream infections in high-income countries, with an estimated incidence of 48 cases per 100,000 person each year (Bonten et al., 2021; Maldonado et al., 2024). A study on early-onset neonatal sepsis involving 235 cases showed that the most frequent pathogen was *E. coli* (36.6%), which was associated with a higher incidence of mortality (Song et al., 2022; Stoll et al., 2020).

Sepsis has been defined as a dysregulated host immune response to infections, leading to a life-threatening organ dysfunction (Song et al., 2022). Sepsis is considered a major cause of mortality worldwide, being responsible for nearly 20% of global deaths (Maldonado et al., 2024).

Several multidrug-resistant (MDR) *E. coli* strains have been reported from sepsis patients (Kumar et al., 2024; Gashaw et al., 2024; Madrazo et al., 2023), making sepsis treatment challenging. Understanding the resistance mechanisms of sepsis-causing MDR pathogens enables the selection of effective antimicrobial agents, thereby improving clinical outcomes and minimising treatment failure. This approach also supports antimicrobial stewardship by reducing reliance on empiric antibiotic therapy and limiting the emergence of resistance.

This study describes the application of *in vitro* phenotypic and genomic approaches for the identification and antimicrobial characterization of a novel MDR *E. coli* strain isolated from the blood of a sepsis patient.

## METHODS

### Bacterial isolation and MALDI-TOF Identification

Following standard microbiological procedures for blood culture processing (CLSI, 2022) a bacterial isolate was obtained from blood sample of a sepsis patient a St. George’s Hospital, London, United Kingdom. A fully automated microbial mass spectrometry detection system (Autof MS1600 MALDI-TOF, Autobio Labtec Instrument Co., Ltd., China) was used in the initial identification of the isolate. The equipment comprised of sample target plate, workstation, (microcomputer, LCD, capture card) instrument (matrix-assisted laser desorption/ionization time of flight detector, vertical ion flight tube, vacuum system), software and database. Sample preparation was carried out according to the Direct Coating Method described in the manual for MALDI-TOF analysis (available at https://chirus.com/autof-ms1600/). First, pure colonies of the bacterial isolates were aseptically transferred to the respective spots on the target plate. A volume of 1 μL of matrix solution (provided by the manufacturer) was then added to each spot containing the sample, and the target plate was placed on a heating block set at 40°C for 1 minute to dry. The target plate was then placed in a holder, which was inserted into the instrument and the analysis was performed. The generated mass spectra were automatically compared with the instrument’s reference database of known spectra from different bacterial species. Based on the closest match from the database, the system provided an identification of the bacterial species.

### Antibiotic Susceptibility Test (AST)

AST of the isolate was carried out using the disk diffusion method described by Chukwunwejim et al. (2018). A volume of 80 mL of molten Mueller Hinton (MH) agar (Sigma-Aldrich, India) was poured into 150 x 20 mm Petri plates and left to solidify. The isolate was grown overnight in MH broth. Using a sterile swab stick, standardized broth culture (0.5 McFarland turbidity standard; OD600 ∼ 0.08 - 0.1) of the isolate was swabbed onto the agar plates. Antibiotic discs [amoxicillin (25 μg), streptomycin (300 μg), cefotaxime (5 μg), oxytetracycline (30 μg), tobramycin (10 μg), and ciprofloxacin (10 μg) (Oxoid, UK)] were placed on the agar surface at a considerable distance between each other. The plates were left to stand for 15 min to allow for the antibiotics to diffuse, after which the plates were incubated at 37°C for 24 h. At the end of incubation, the plates were observed for inhibition zone diameters (IZDs), and these were measured in mm using a meter rule. *E. coli* (ATCC 25922), a non-MDR (antibiotic-susceptible) strain, was used as a control. The experiment was conducted in triplicate and the mean IZDs were calculated and recorded.

### MIC Determination

The minimum inhibitory concentrations (MICs) of several antibiotics [ciprofloxacin (Merck, USA), gentamicin sulfate salt (Sigma-Aldrich, Germany), and neomycin sulphate (Apollo Scientific, UK)] on the isolate were determined by broth microdilution method in cation adjusted Mueller Hinton broth (CAMHB) (Sigma-Aldrich, India) using a final bacterial inoculum of 1x10^6^ CFU/ml (Oyama et al., 2022). The antibiotics, dissolved in sterile distilled water, were added to the bacterial culture and incubated overnight (18–24 h) at 37 °C. After incubation, the wells were visually examined for the presence or absence of growth, indicated by the turbidity of the medium. *E. coli* (ATCC 25922), a non-MDR strain, was used as a control. The MICs of the antibiotics were recorded as the lowest concentration that inhibited growth (prevented visible growth of bacteria).

### Genomic DNA extraction, sequencing, and assembly

Genomic DNA extraction, hybrid (Illumina & Nanopore) sequencing, and assembly of the bacterial isolate was carried out by a commercial service provider, MicrobesNG (Birmingham, UK), according to protocols available at https://microbesng.com/documents/methods/. The bacterial DNA was subjected to HYBRID (Illumina & Nanopore) sequencing which combines data from the latest R10.4.1 chemistry from Oxford Nanopore Technologies with 2 x 250 bp Illumina reads to provide high quality assemblies of bacterial genomes. Hybrid assembly was performed using Unicycler (version 0.4.0; Wick et al., 2017). QUAST on Galaxy https://usegalaxy.org/; Mikheenko et al. 2018; Gurevich et al., 2013) was used to assess the quality of the genome assembly.

### Genome component prediction and gene annotation

The hybrid assembled genome was annotated using Prokka (version 1.14.6) available on Galaxy (https://usegalaxy.org/) (Seemann, 2014). PlasmidFinder (Galaxy Version 2.1.6+galaxy1) and MOB-Recon (Galaxy Version 3.0.3+galaxy0), both available on Galaxy, were also used to assess the properties of the contigs and determine whether they were chromosomes or plasmids, as well as to identify plasmid type and circularity (Carattoli et al., 2014; Robertson and Nash, 2018). “Extractseq” on Galaxy was used to extract the 16S rRNA sequence from the bacterial genome for species identification via BLAST searches on NCBI, Greengenes and SILVA databases. The GC content of the complete genome and the contigs were calculated using the GC Content Calculator available at https://jamiemcgowan.ie/bioinf/gc_content.html. Antimicrobial resistance genes (ARGs) were identified in the bacterial genome using tools available on Galaxy: AMRFinderPlus (Galaxy Version 3.12.8+galaxy0), an NCBI antimicrobial resistance gene (ARG) finder, and starAMR (Galaxy Version 0.11.0+galaxy0), which searched against the ResFinder, PlasmidFinder, and PointFinder databases. Additionally, the Resistance Gene Identifier (RGI; version: 6.0.5) from the Comprehensive Antibiotic Resistance Database (CARD; version: 4.0.1), available at https://card.mcmaster.ca/analyze/rgi (Alcock et al., 2023), was used to identify the ARGs in the bacterial genome. Functional annotation of the bacterial genome was performed using the eggNOG-mapper tool (version: 2.1.12) with the eggNOG 5.0 database, available at http://eggnog-mapper.embl.de/ (Cantalapiedra et al., 2021).

### Isolate identity confirmation by 16S taxonomy

To confirm the identity of the isolate, its 16S rRNA gene (in FASTA format) was analysed using the standard and rRNA/ITS databases of the NCBI BLAST search tool, available at https://blast.ncbi.nlm.nih.gov/Blast.cgi (Altschul et al., 1990). Additionally, the 16S rRNA sequence was queried against the SILVA 138.2 database (https://www.arb-silva.de/) using the SILVA ACT (Alignment, Classification, and Tree) tool, which allows for sequence upload and comparison against their curated ribosomal RNA database (Pruesse et al., 2012). The appropriate search parameters, such as targeting the 16S rRNA gene for bacteria, were selected, and the search was executed. The 16S rRNA sequence was also analysed using the Greengenes2 2024.09 reference database (McDonald et al., 2024), available at https://github.com/Greengenes2/greengenes2. This database is designed to classify microbial 16S rRNA sequences based on taxonomic hierarchy by comparing the query sequence against a curated reference set to identify close matches and assign taxonomy.

Sourmash (version 4.8.10) was used to cluster the isolate with closely related species identified in the 16S rRNA NCBI BLAST search, aiming to clarify its taxonomic species-specific identity by comparing it to various closely related species’ genomes found in the BLAST search (Brown and Irber, 2016). To avoid misclassification due to plasmid presence, only the bacterial chromosome was analysed. A total of 33 additional genomes of closely related species, identified in the 16S rRNA BLAST search, were downloaded from the NCBI database, with two genomes for each bacterial species identified as close hits in the BLAST search (see Supplementary Material: Figure S7). These genomes, along with the isolate’s chromosome were analysed in Sourmash using the Jaccard Index with a k-mer size of 31. To further confirm the results, a second Sourmash plot was generated using the isolate’s chromosome and an additional 1,594 bacterial genomes downloaded from the Bacterial and Viral Bioinformatics Resource Center (BV-BRC) database, available at https://www.bv-brc.org/ (see *wget*.*sh* and *genome_list*.*txt* at https://github.com/LucyDillon/MDR_isolate for details). The estimated average nucleotide identity (ANI) and Jaccard Index of the 1,595 isolates were compared using a k-mer size of 31.

### Phylogenetic relatedness of isolate

A pangenome analysis was conducted to produce a core genome alignment of the closely related species based on the 16S rRNA BLAST search. To generate the core gene alignment, the genomes were first annotated using Prokka (version 1.14.5; Seemann, 2014). Due to the large number of genomes, this step was performed via high-performance computing (HPC). Roary (version 3.13.0) was then used for the pangenome analysis (Page et al., 2015) (see *Roary_core gene*.*sh* at https://github.com/LucyDillon/MDR_isolate for parameter details). Roary produces a default Newick file via FastTree (Price et al., 2009), which was visualized using iTOL (Letunic and Bork, 2024). Additionally, another phylogenetic tree was generated from the core gene alignment using IQ-TREE (version 2; Minh et al., 2020), and the resulting tree was visualized in iTOL (Nguyen et al., 2015) (see *iq_tree*.*sh* at https://github.com/LucyDillon/MDR_isolate for parameter details).

### Biochemical differentiation and phenotypic confirmation of isolate

To confirm the phenotypic and biochemical similarity or dissimilarity of the isolate to *E. coli* or *E. fergusonii*, and to ascertain if the isolate is a *Shigella* species, the isolate and reference strains of *E. coli* (ATCC 25922) and *E. fergusonii* (ATCC 35469) were streaked on Coliform *ChromoSelect* Agar (CCA; Sigma-Aldrich, Germany) and incubated overnight at 37°C. CCA is a selective medium used for the detection of *E. coli* and total coliforms in water and food samples. The chromogenic mixture contains two chromogenic substrates: Salmon-GAL and X-glucuronide. The enzyme β-D-galactosidase, produced by coliforms (including *E. fergusonii* strains), cleaves Salmon-GAL, resulting in a colour change from salmon to red in the coliform colonies. *E. coli* produces dark blue to violet colonies on CCA because it cleaves both Salmon-GAL and X-glucuronide using β-D-galactosidase and β-D-glucuronidase, respectively, and exhibits a positive indole reaction, as the medium contains tryptophan. *Shigella* spp., being non-coliform, appear colourless on CCA, as they are negative for Salmon-GAL, X-Gluguronide, and indole (Sigma-Aldrich, 2024).

## RESULTS

### Preliminary Strain identification and Phenotypic Resistance Profiling

MALDI-TOF analysis identified the isolate as *Escherichia coli* (see Supplementary Material: Figure S1). It was then ascribed the strain-level identity *E. coli* 266631E. Resistance to several commonly used antibiotics was confirmed through disk diffusion testing and MIC determination (Table 1). Compared to the non-MDR reference strain *E. coli* strain (ATCC 25922), the isolate exhibited resistance to amoxicillin, cefotaxime, tobramycin and ciprofloxacin, with inhibition zone diameters (IZD) of 0 mm for all except for tobramycin, which had an IZD of 11 mm. Additionally, MIC determination by broth dilution assay revealed that the isolate was highly resistant to ciprofloxacin and gentamicin, with MIC values of 31.25 and 500 μg/mL, respectively, compared to the non-MDR control strain.

**Table 1.** Phenotypic Resistance Profile of the Bacterial Isolate

| Antibiotics | Inhibition Zone Diameter (mm) |  |
| --- | --- | --- |
|  | *E. coli* 266631E | *E. coli* (ATCC 25922) |
| Amoxicillin (25 µg) | 0 | 20 |
| Streptomycin (300 µg) | 22 | 22 |
| Cefotaxime (5 µg) | 0 | 28 |
| Oxytetracycline (30 µg) | 23 | 26 |
| Tobramycin (10 µg) | 11 | 20 |
| Ciprofloxacin (10 µg) | 0 | 38 |
| MIC (µg/mL) |  |  |
| Ciprofloxacin | 31.25 | 0.015 |
| Gentamicin | 500 | 3.9 |
| Neomycin | 3.9 | 3.9 |

### Genomic Insights into Antimicrobial Resistance and Functional Gene Annotations

The complete genome sequence of *E. coli* 266631E is 5,454,694 bp in size and consists of 10 contigs. Contig 1 represents the chromosome (5,019,723 bp), while the remaining nine contigs include those containing plasmids such as IncF-type (contigs 3 and 4), Col156 (contigs 5), and ColMG828 (contig 8) (Table 2; Supplementary Material: Figures S2 – S5). ARGs identified using AMRFinderPlus, starAMR and RGI revealed a collection of ARGs including *aadA5*, CTX-M-15, *dfrA17, mphA, qacE, sul1*, and TEM-1 as outlined in Table 2 below.

**Table 2.** Genome assembly information and ARG carriage in *E. coli* 266631E

| Contig | Total length (bp) | GC content | Plasmid type [As determined by starAMR (†), Plasmid finder (‡), and MOB-Recon (§)] | Circularity status [As determined by Unicycler (†), Plasmid finder (‡), and MOB-Recon (§)] | Predicted ARGs [As determined by AMRFinderPlus (%Identity: 100%)] | Predicted ARGs [As determined by starAMR (%Identity: ≥99%)] | Predicted ARGs [As determined by RGI (%Identity: ≥99%)] | NCBI Genome Accession Number* |
| --- | --- | --- | --- | --- | --- | --- | --- | --- |
| Contig 1 (Chromosome) | 5019723 | 50.77% | - | - | CTX-M-15 (2 copies) | CTX-M-15 (2 copies) | *acrA*, *acrAB-TolC* (with *acrR* mutation), *acrB*, *acrD*, *acrE*, *acrF*, *bacA*, *cpxA*, *CRP*, *CTX-M-15* (2 copies), *emrA*, *emrB*, *emrR*, *emrY*, *evgA*, *gadW*, *H-NS*, *kdpE*, *leuO*, *marA*, *mdtB*, *mdtE*, *mdtF*, *mdtG*, *mdtH*, *msbA*, *PmrF*, *soxR*, *soxS*, *TolC*, *ugd*, *Yojl* | CP183489 |
| Contig 2 | 218237 | 50.13% | - | Circular<sup>†§</sup> | None | None | None | CP183490 |
| Contig 3 | 126566 | 52.54% | IncF-type<sup>†‡§</sup> | Circular<sup>†‡§</sup> | *aac(3)-II*, *aadA5*, *CTX-M-15*, *dfrA17*, *mph(A)*, *qacE*, *sul1*, TEM-1B | *aac(3)-II*, *aadA5*, *CTX-M-15*, *dfrA17*, *mph(A)*, *qacE*, *sul1*, TEM-1B | *aadA5*, *CTX-M-15*, *dfrA17*, *mphA*, *Mrx*, *qacEdelta1*, *sul1*, TEM-1 | CP183491 |
| Contig 4 | 74099 | 51.25% | IncF-type<sup>†‡§</sup> | Circular<sup>†‡§</sup> | None | None | None | CP183492 |
| Contig 5 | 6078 | 49.03% | Col156<sup>‡§</sup> | Circular<sup>†‡§</sup> | None | None | None | CP183493 |
| Contig 6 | 4600 | 45.74% | - | - | None | None | None | CP183494 |
| Contig 7 | 1709 | 54.13% | - | - | None | None | None | CP183495 |
| Contig 8 | 1552 | 51.87% | ColMG828<sup>§</sup> | Circular<sup>†§</sup> | None | None | None | CP183496 |
| Contig 9 | 1307 | 47.28% | - | Circular<sup>†§</sup> | None | None | None | CP183497 |
| Contig 10 | 823 | 50.67% | - | - | None | None | None | CP183498 |

\*The isolate's complete genome, including the chromosome, plasmids, and the other contigs, was deposited in the NCBI Genome database under the accession numbers CP183489 – CP183498.

Using the eggNOG-mapper tool, the functional gene families across the bacterial isolate were identified. The COG (clusters of orthologous groups) categories that constituted the largest gene families within the bacterial genome were M (Cell wall/membrane/envelope biogenesis), P (Inorganic ion transport and metabolism), and K (Transcription) (Figure 1). The majority of the RGI predicted ARGs identified in the bacterial chromosome were grouped into their respective COG categories, as indicated in the eggNOG-mapper output file (Table 3).

**Table 3.** Classification of the identified ARGs into functional gene families by COG categories

| COG category | RGI predicted genes |  |  |
| --- | --- | --- | --- |
|  | ARGs | AMR mechanism | Target Drug Class |
| M (Cell wall/membrane/envelope biogenesis) | *acrA* | Antibiotic target alteration, antibiotic efflux | Fluoroquinolones, cephalosporins, glycylicycline, penams, tetracyclines, rifamycins, phenicols, disinfecting agents, antiseptics |
|  | *tolC* | Antibiotic efflux | Macrolides, fluoroquinolones, aminoglycosides, carbapenem, cephalosporins, glycylicycline, cephamycins, penams, tetracyclines, peptide antibiotic, aminocoumarins, rifamycins, phenicols, penems, disinfecting agents and antiseptics |
|  | *mdtE* | Antibiotic efflux | Macrolides, fluoroquinolones, penam |
| P (Inorganic ion transport and metabolism) | *yojI* | Antibiotic efflux | Peptide antibiotic |
| K (Transcription) | *soxS* | Antibiotic target alteration, antibiotic efflux, reduced permeability to antibiotic | Fluoroquinolones, monobactams, carbapenems, cephalosporins, glycylicyclines, cephamycins, penam, tetracyclines, rifamycins, phenicols, penems, disinfecting agents, antiseptics |
|  | *soxR* | Antibiotic target alteration, antibiotic efflux | Fluoroquinolones, cephalosporins, glycylicyclines, penams, tetracyclines, rifamycins, phenicols, disinfecting agents and antiseptics |
|  | *crp* | Antibiotic efflux | Macrolides, fluoroquinolones, penams |
|  | *evgA* | Antibiotic efflux | Macrolides, fluoroquinolones, penams, tetracyclines |
|  | *kdpE* | Antibiotic efflux | Aminoglycosides |
|  | *H-NS* | Antibiotic efflux | Macrolides, fluoroquinolones, cephalosporins, cephamycins, penams, tetracyclines |
|  | *gadW* | Antibiotic efflux | Macrolides, fluoroquinolones, penam |
|  | *acrAB-TolC*<br>(with *acrR* mutation) | Antibiotic target alteration, antibiotic efflux | Fluoroquinolones, cephalosporins, glycylicycline, penams, tetracyclines, rifamycins, phenicols, disinfecting agents, antiseptics |
|  | *marA* | Antibiotic efflux, reduced permeability to antibiotics | Peptide antibiotics |
|  | *leuO* | Antibiotic efflux | Nucleosides, disinfecting agents, antiseptics |
| U (Intracellular trafficking, secretion, and vesicular transport) | *acrD* | Antibiotic efflux | Aminoglycosides |
|  | *emrB* | Antibiotic efflux | Fluoroquinolones |
| V (Defence mechanisms) | *acrB* | Antibiotic efflux | Fluoroquinolones, cephalosporins, glycylicyclines, penams, tetracyclines, rifamycin, phenicol, disinfecting agents, antiseptics |
|  | *mdtB* | Antibiotic efflux | Aminocoumarins |
|  | *mdtF* | Antibiotic efflux | Macrolides, fluoroquinolones, penams |
|  | *msbA* | Antibiotic efflux | Nitroimidazoles |
|  | *emrA* | Antibiotic efflux | Fluoroquinolones |
| T (Signal transduction mechanisms) | *cpxA* | Antibiotic efflux | Aminoglycosides, aminocoumarins |
| C (Energy production and conversion) | *ugD* | Antibiotic target alteration | Peptide antibiotics |

**Figure 1.**
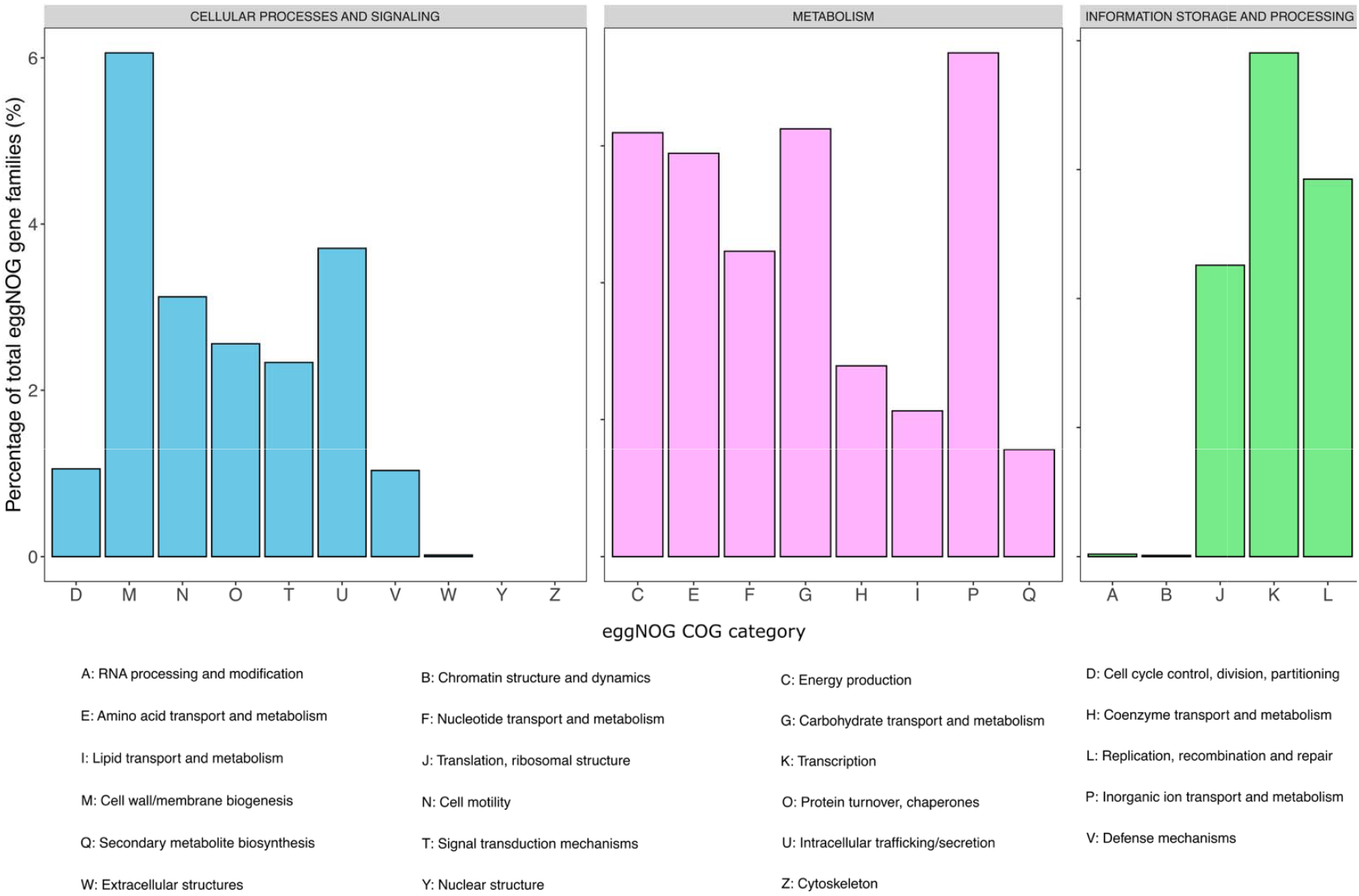
The distribution of eggNOG gene families across *E. coli* 266631E genome

### Taxonomy

#### Query of the isolate’s 16S rRNA sequence against NCBI, Greengenes2, and SILVA Databases

A BLAST search of the bacterial 16S rRNA sequence in the NCBI standard nucleotide databases identified the isolate as *E. coli* with 100% identity (Supplementary Material: Figure S6). However, a BLAST search of the bacterial 16S rRNA sequence in the NCBI rRNA/ITS databases showed the highest percent identity to two *Shigella* species – *S. sonnei* (99.80%) and *S. flexneri* (99.80%), followed by two *E. fergusonii* strains (99.66% and 99.54%, respectively) and then *E. coli* (99.52%) (Supplementary Material: Figure S7). Furthermore, querying the bacterial 16S rRNA sequence against the Greengenes2 database revealed 100% identity matches for 1,487 out of 6,066 bacterial strains in the database. Among these 1,487 strains, the highest species-specific match was for *E. fergusonii* with 130 matches, followed by 30 matches for *E. coli*, and 10 matches for *Shigella* species. Interestingly, most of the 100% matches were for non-species-specific bacteria in the *Escherichia* genus (Supplementary Material: Figure S8). Finally, a sequence alignment of the bacterial 16S rRNA sequence against the SILVA reference database yielded a non-species-specific and non-genus-specific taxonomic classification: ‘Escherichia-Shigella’ (Supplementary Material: Figure S9).

#### Phylogeny of the bacterial isolate

As the query of the isolate’s 16S rRNA sequence against NCBI, Greengenes2, and SILVA databases did not align in terms of taxonomy, Sourmash was used to cluster the isolate (labelled as ‘‘unknown isolate’’; Figure 2) with additional genomes. A total of 33 additional genomes of closely related species, identified in the 16S rRNA NCBI BLAST search, were downloaded, with two genomes for each bacterial species identified as close hits in the BLAST search (i.e., *E. coli, E. fergusonii* and *Shigella* spp.). Sourmash Jaccard index analysis showed that the closest genome was *E. coli*; however, several *Shigella* spp. clustered closer to the isolate than the second *E. coli* genome, making it difficult to definitively identify the isolate as *E. coli*. Since the initial Sourmash analysis was not entirely conclusive—possibly due to the small number of genomes analysed—and to enable a more comprehensive Sourmash clustering analysis, an additional 1,594 genomes were sourced from the BV-BRC database (a comprehensive, curated repository of bacterial and viral genomes that provides access to a vast collection of high-quality, annotated reference sequences). Both the ANI plot (Supplementary Material: Figure S10) and the Jaccard Sourmash plot (Supplementary Material: Figure S11) showed that the isolate clustered among several *Shigella* and *Escherichia* species.

**Figure 2.**
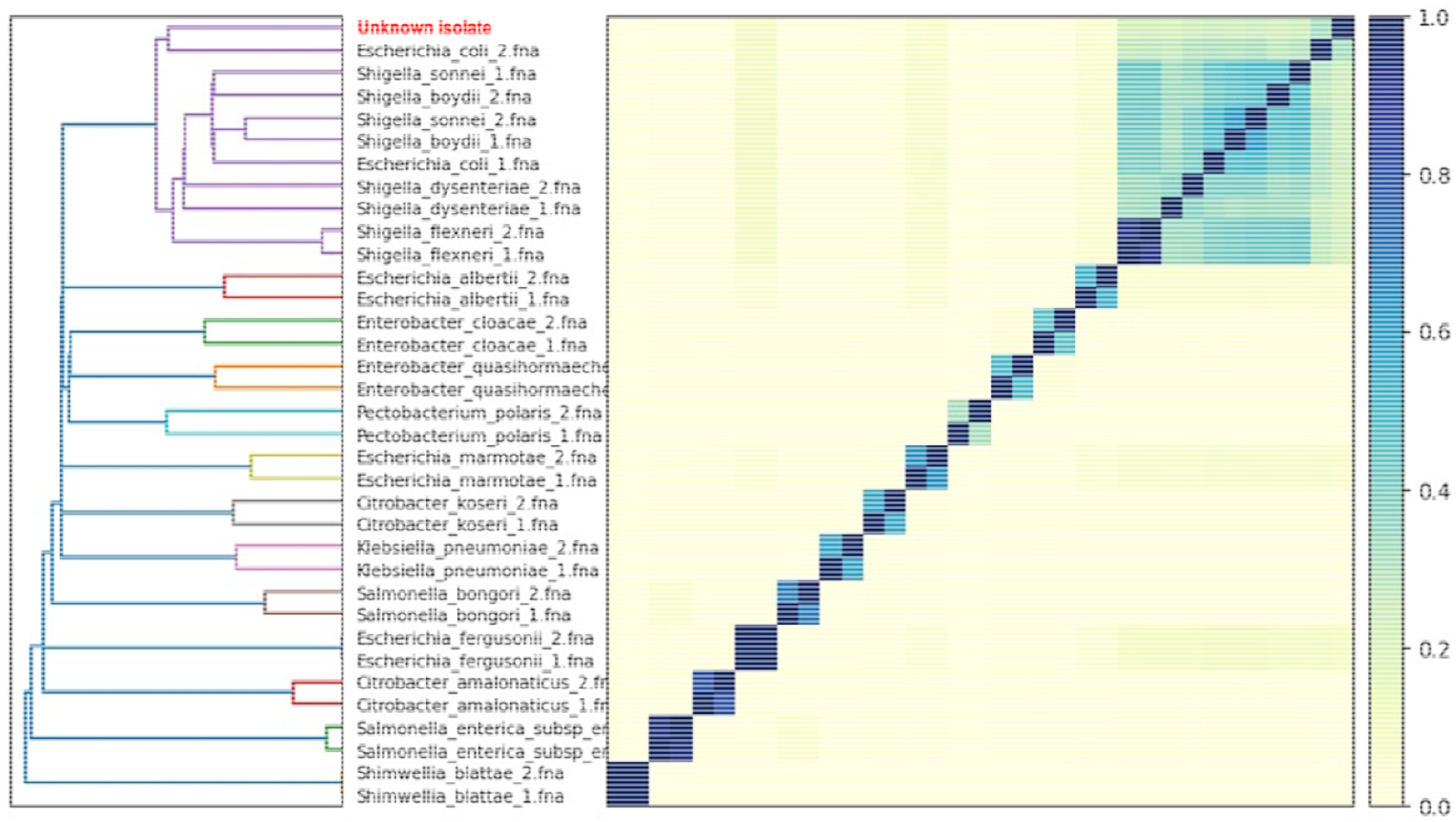
Sourmash plot of 34 genomes (the isolate’s chromosome plus 33 additional genomes of closely related species identified in the 16S rRNA BLAST search). The plot showed that the closest genome to the isolate is *E. coli*. However, several *Shigella* spp. also clustered closer to the isolate than the second *E. coli* genome, complicating the definitive identification of the isolate as *E. coli*.

Due to the ambiguous results of the Sourmash analysis regarding the species-specific identification of the isolate, a pangenome analysis was conducted to produce a core genome alignment of closely related species identified by the 16S rRNA BLAST search. The 1,595 isolates from the Sourmash analysis were used to construct the phylogeny tree. Roary was employed to generate a core gene alignment and create a Newick file with FastTree, which was visualized in iTOL (Supplementary Material: Figure S12). The isolate clustered with a mix of *Shigella* and *Escherichia* species, consistent with the patterns observed in the Sourmash plots.

The phylogenetic tree produced by FastTree did not provide a definitive or conclusive taxonomy (i.e., species-specific identification) for the isolate. Therefore, a more rigorous phylogenetic tool, IQ-TREE, was used to generate another tree from the core gene alignment. IQ-TREE is known for its advanced modelling of evolutionary processes, stronger statistical support for tree branches, and greater reliability for species-level resolution— particularly in closely related taxa such as *Escherichia* and *Shigella* (Nguyen et al., 2015; The et al., 2021; Minh et al., 2020). The resulting tree was visualized in iTOL, and branches from unrelated clades were removed to provide a clearer visualization of the isolate and surrounding bacterial species. The IQ-TREE plot yielded the same result as the FastTree plot, as well as the Sourmash plots, showing the isolate clustering with *Escherichia* and *Shigella* species (Figure 3).

**Figure 3.**
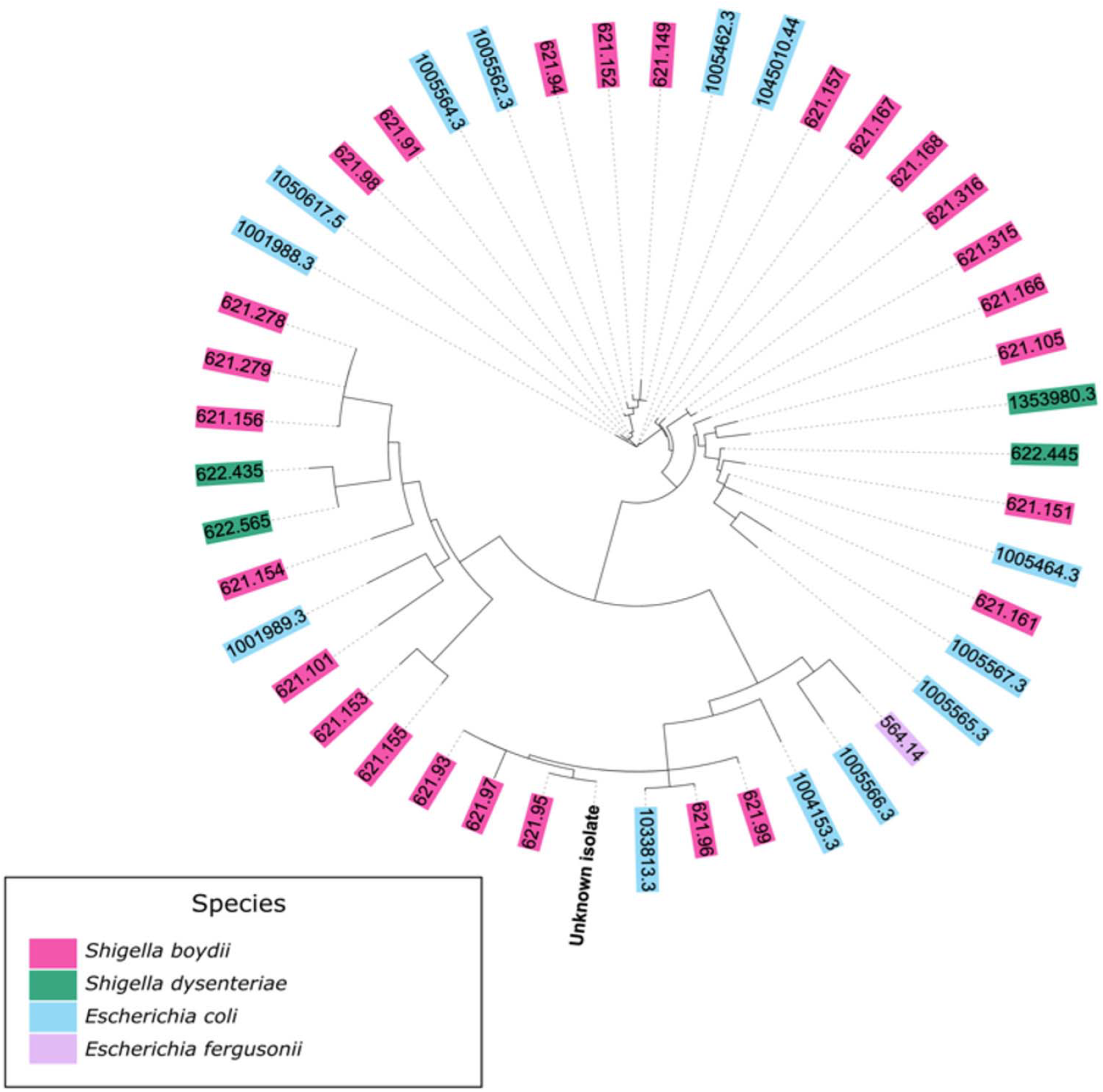
Phylogeny of the isolate using IQ-TREE. The tree is a reduced representation of 1,595 isolates; and clades have been removed to be able to visualise the isolate (labelled as ‘‘unknown isolate’’) and surrounding genomes.

#### Biochemical differentiation of isolate

By applying a simple phenotypic, biochemical, and cultural approach to confirm the identity of the isolate or distinguish it from *E. coli, E. fergusonii*, or *Shigella* species, the isolate and typed strains of *E. coli* (ATCC 25922) and *E. fergusonii* (ATCC 35469) were cultivated on CCA, a chromogenic selective agar that differentiates *E. coli* from other coliforms. The isolate, like the typed *E. coli* strain (ATCC 25922), produced characteristic dark blue to violet-coloured colonies on CCA due to cleavage of both Salmon-GAL and X-glucuronide in the selective chromogenic culture medium. In contrast, *E. fergusonii* (ATCC 35469) produced pink colonies (see Figure 4). Based on the biochemical/chromogenic characteristics, the isolate can be identified as an *E. coli* strain and is clearly distinguishable from *E. fergusonii*, and from *Shigella* species, which are known to produce colourless colonies on CCA (Sigma-Aldrich, 2024).

**Figure 4.**
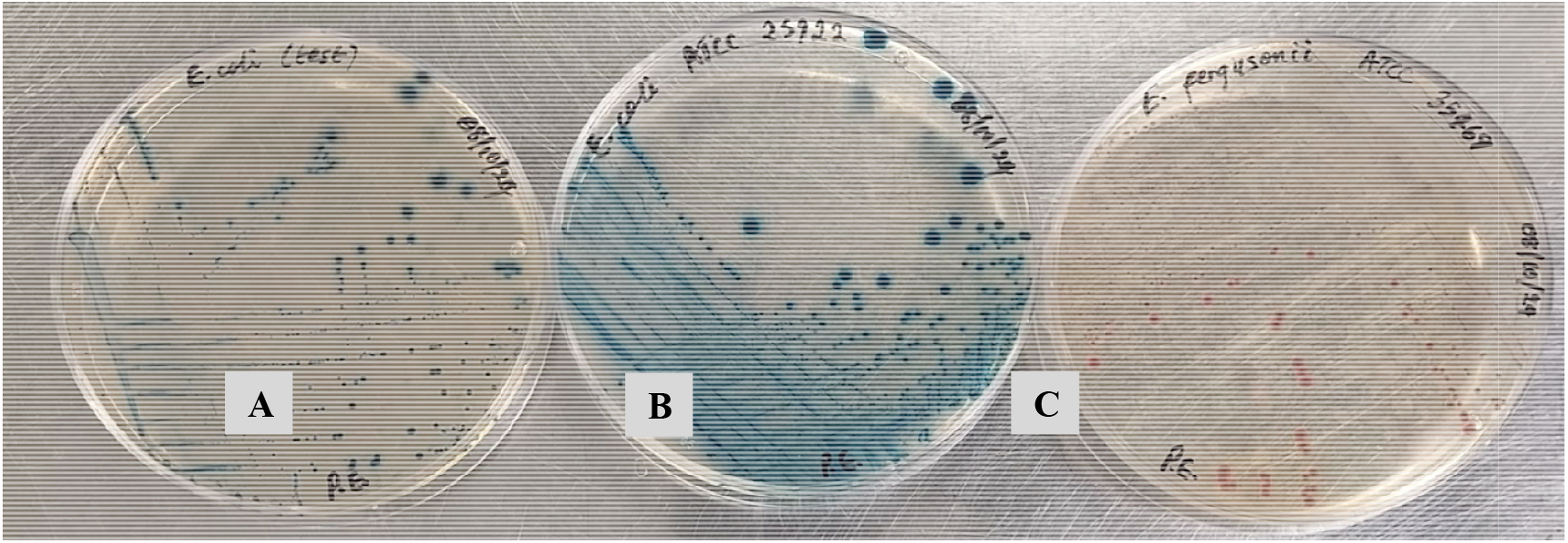
The colony characteristics of the isolate (A) on CCA showed dark blue to violet pigmentation, typical of *E. coli*, as confirmed by the dark blue to violet colonies of the reference *E. coli* strain (ATCC 25922) (B). The isolate is clearly distinguishable from the pink colonies of *E. fergusonii* (C) and from *Shigella* species, which are known to produce colourless colonies on CCA.

## DISCUSSION

*E. coli*, one of the most extensively studied microorganisms in the world, is an opportunistic pathogen comprising strains that typically exist as harmless bacteria, forming a normal part of the gastrointestinal flora in many warm-blooded animals and reptiles. However, several highly adapted commensal clones can acquire specific virulence attributes, becoming highly pathogenic and capable of causing life-threatening infections, such as sepsis (Peng et al., 2024; Geurtsen et al., 2022).

*E. coli* sepsis, an invasive ExPEC disease, is a systemic infectious disease characterized by severe fever, chills, tachycardia, hypotension, and multiple organ dysfunction syndrome, which are typical clinical symptoms (Shao et al., 2023; Doua et al., 2023). ExPEC is now the leading cause of invasive bacterial disease globally, surpassing pathogens like *Staphylococcus aureus* and *Streptococcus pneumoniae. E. coli* is responsible for over 40% of bloodstream infections, with a case-fatality rate (CFR) of 12.4%. Sepsis, often caused by *E. coli*, accounted for 11 million deaths in 2017 from nearly 49 million cases globally (Doua et al., 2023; Pérez-Crespo et al., 2021; Bonten et al., 2021; Buetti et al., 2016).

Early sepsis diagnosis and timely and appropriate antibiotic therapy are crucial in preventing the progression of sepsis to septic shock or multiorgan failure. However, treatment can be delayed due to misdiagnosis or lack of knowledge about the pathogen (Doua et al., 2023). Antimicrobial resistance in ExPEC, particularly concerning MDR strains that cause sepsis, such as extended-spectrum β-lactamase (ESBL)-producing *E. coli*, poses a major threat to effective treatment. This underscores the need to understand resistance mechanisms to guide antibiotic selection and improve survival rates (Kumar et al., 2024; Gashaw et al., 2024; Doua et al., 2023; Madrazo et al., 2023).

This study reports the isolation and identification of an MDR *E. coli* strain (*E. coli* 266631E) from the blood of a sepsis patient, as well as the characterization of the genetic determinants of AMR in the bacterial isolate.

### Phenotypic AMR Screening and Antimicrobial Resistance Gene (ARG) Analysis

Both phenotypic AMR screening (disk diffusion testing and broth microdilution MIC determination) and ARG analysis demonstrated the multi-drug resistance of the bacterial isolate. Results from the AST and MIC determination experiments show that *E. coli* 266631E was resistant to multiple antibiotics, including amoxicillin, cefotaxime, tobramycin, ciprofloxacin, and gentamicin (Table 1). WGS and analysis of the genetic determinats of AMR in the isolate reveal a multitude of ARGs responsible for diverse mechanisms of resistance to a vast range of antimicrobials across different classes.

Genomic analysis of *E. coli* 266631E revealed genes encoding for resistance to clinically important antibiotics, including those from the macrolide, fluoroquinolone, aminoglycoside, cephamycin, glycylcycline, tetracycline, peptide antibiotic, aminocoumarin, rifamycin, phenicol, cephalosporin, penam, and penem classes (Tables 2 and 3). Multiple ARGs with diverse mechanisms of action and responsible for resistance to a wide range of antibiotics from different classes, were detected in the bacterial chromosome using RGI (Table 2). However, only *CTX-M-15* was identified in the chromosome using the AMRFinderPlus and starAMR databases. Compared to AMRFinderPlus and starAMR, RGI appears to be more robust in identifying ARGs in the bacterial chromosome. Outside the chromosome, ARGs were detected only in contig 3, identified as an IncF-type plasmid. In this plasmid, except for the *Mrx* gene, which was detected only by RGI, and *aac(3)-II*, detected only by AMRFinderPlus and starAMR, all three AMR prediction tools (AMRFinderPlus, starAMR, and RGI) equally detected seven similar ARGs: *aadA5*, CTX-M-15, *dfrA17, mphA, qacE, sul1*, and TEM-1.

The CTX-M-15, a key ARG of public health concern and one of the most common and influential genotypes of the *bla*_CTX-Ms_ family, is widely present in *E. coli* (Yu et al., 2024; Rangama et al., 2021). This gene, identified both in the bacterial chromosome (with duplicate copies in different loci) and the IncF-type plasmid identified in contig 3 (Table 2), is known to encode for ESBL, which targets beta-lactam antibiotics, including third-generation cephalosporins (such as cefepime, cefotaxime, ceftriaxone) and penicillins (Rossolini et al., 2008; Livermore and Hawkey, 2005). Resistance to carbapenems has also been observed in CTX-M-15-producing bacteria (Elliott et al., 2006; Livermore and Hawkey, 2005). As observed in the *E. coli* strain isolated in this study, this MDR gene is frequently carried on plasmids, facilitating its rapid spread between different bacterial species (Yu et al., 2024; Grevskott et al., 2024).

*E. coli* 266631E exhibits a critical convergence of chromosomal and plasmid-mediated resistance mechanisms frequently observed in bloodstream infections (Kirtikliene et al., 2022; Goswami et al., 2020). Its unusually high chromosomal ARG load suggests a significant potential to pose treatment challenges. The isolate’s high-level quinolone resistance is evidenced by the presence of multiple quinolone-resistance genes within the bacterial genome. Most of the identified chromosomal ARGs are linked to fluoroquinolone resistance, including *acrA, acrB, crp, evgA, gadW, H-NS, acrAB-TolC* (with an *acrR* mutation), *emrA, emrB, mdtE, mdtF, soxR, soxS*, and *tolC*. These genes confer resistance through diverse mechanisms, including antibiotic efflux, target alteration, and reduced permeability. The abundance of these chromosomal fluoroquinolone-resistance genes aligns with established efflux- and regulator-mediated resistance mechanisms in clinical *E. coli* isolates (Esmaeel et al., 2020; Amin et al., 2021). Studies have shown that MDR E. coli often amplify efflux pump genes (e.g., *acrAB-TolC, emr, mdt*) as a primary defense against fluoroquinolones and other antibiotics (Nasrollahian et al., 2024; Altayb et al., 2022; Rozwadowski and Gawel, 2022).

Several aminoglycoside resistance genes were identified in the chromosome (*tolC, kdpE, acrD, cpxA*) and in the IncF-type plasmid on contig 3 (*aadA5 and aac(3)-II)* (Table 2; Supplementary Material: Figures S2–S5). IncF-type plasmids, common in hospital-associated bloodstream isolates, frequently encode aminoglycoside-modifying enzymes and ESBLs (Mangroliya et al., 2025; Liu et al., 2022; Yang et al., 2015). The absence of quinolone resistance genes in this plasmid suggests that quinolone resistance in *E. coli* 266631E is chromosomally mediated. The presence of fluoroquinolone and aminoglycoside resistance genes in both the chromosome and plasmid correlates with the high resistance levels observed in the AST (Table 1). Compared to the reference strain *E. coli* (ATCC 25922), which falls within EUCAST’s susceptibility range (EUCAST, 2023), *E. coli* 266631E displays strong resistance to ciprofloxacin (10 µg disc, 0 mm IZD; MIC: 31.25 µg/mL), gentamicin (MIC: 500 µg/mL), and reduced susceptibility to tobramycin (10 µg disc, 11 mm IZD).

Functional classification of the chromosomal ARGs using eggNOG-mapper revealed that most fall into COG categories M (cell wall/membrane/envelope biogenesis), K (transcription), and V (defence mechanisms) (Table 3, Figure 1). Table 3 shows that ARGs within each category employ diverse resistance mechanisms, and not just single-gene effects. Notably, genes with similar resistance functions, such as those mediating antibiotic efflux, are distributed across multiple COG categories, including M, P, K, U, V, and T. This distribution indicates a complex and versatile resistance mechanism that enhances the bacterium’s ability to withstand various antibiotics. The genetic diversity and functional spread of these ARGs underscore the clinical significance of the strain, highlighting its potential to pose treatment challenges.

The detection of such a highly resistant *E. coli* strain in a hospitalized sepsis patient is concerning. Hospital environments, particularly intensive care units (ICUs), are hotspots for MDR pathogens, where plasmid-mediated ARGs can transfer across strains (Mangroliya et al., 2025). The combination of chromosomal and plasmid-borne resistance significantly complicates treatment and heightens the risk of nosocomial transmission.

### Taxonomy

An attempt to confirm the identity of the bacterial isolate by querying the bacterial 16S rRNA sequence against the NCBI, Greengenes, and SILVA databases produced variable outcomes that did not align taxonomically. A BLAST search of the bacterial 16S rRNA sequence in the NCBI standard databases identified the isolate as *E. coli* with 100% identity. However, when searched in the NCBI rRNA/ITS databases, the sequence showed the highest identity to *Shigella* species, followed by *E. fergusonii*, and then *E. coli* (Supplementary Material: Figures S6 – S7). Querying the bacterial 16S rRNA sequence against the Greengenes2 database revealed 100% identity matches for *E. fergusonii, E. coli*, and *Shigella* species. Additionally, submitting the 16S rRNA sequence to SILVA yielded a non-species-specific and non-genus-specific taxonomic classification: ‘*Escherichia–Shigella*’ which is consistent with the report by Edgar (2018). Edgar noted that SILVA, which is based on Bergey’s Manual and List of Prokaryotic Names with Standing in Nomenclature (LSPN), addresses the taxonomic overlap between *Escherichia* and *Shigella* by retaining well-established species names, such as *Escherichia coli*, and introducing a combined genus named *Escherichia–Shigella*.

The taxonomic relationship between *Escherichia* and *Shigella* has been a longstanding subject of debate, mainly due to the close genetic and phenotypic similarities between these genera (Sanford et al., 2021; Khot and Fisher, 2013; Lan and Reeves, 2001). Traditional taxonomic methods, such as 16S rRNA sequencing, which has been widely used to classify bacterial species, lack the resolution to effectively distinguish between closely related species such as *Escherichia* and *Shigella* (Lan and Reeves, 2001; Rantakokko-Jalava et al., 2000). The 16S rRNA gene is conserved across most bacteria, making it useful for broader taxonomic classifications (Jo et al., 2017) but insufficient for species-level distinctions in many genera, including *Escherichia* and *Shigella* (Liu et al., 2023; Khot and Fisher, 2013).

Historically, *Shigella* was classified separately from *E. coli* based on clinical, phenotypic, and biochemical differences, especially due to its association with dysentery and other pathogenic traits. There are very few biochemical properties that distinguish *Shigella* from enteroinvasive *E. coli* (EIEC), which is also a major cause of dysentery. In fact, some O-antigens associated with EIEC are identical to those found in *Shigella* spp., and many plasmid-associated virulence determinants are common to both EIEC and *Shigella* (Yang et al., 2005; Lan et al., 2001; Lan and Reeves, 2001; Cheasty and Rowe, 1983). However, advances in genomics, particularly whole genome sequencing, have shown that *Shigella* is genetically nested within the *Escherichia* genus, raising questions about the validity of the historical separation of these genera (Abram et al., 2021; The et al., 2016; Yang et al., 2005). Because they are genetically almost identical to certain strains of *E. coli* but have distinct pathogenic mechanisms that cause specific diseases, particularly dysentery (shigellosis), *Shigella* spp. are considered to be pathotypes of *E. coli* (Pupo et al., 2000, Lan et al., 2004; Lindsey et al., 2017). Despite genomic evidence supporting the integration of *Shigella* into *Escherichia*, the practical implications of such a taxonomic change are significant and could impact clinical diagnostics, epidemiological tracking, and communication among researchers. Therefore, any taxonomic changes must be carefully considered and widely accepted by both the scientific and medical communities to avoid confusion and ensure clarity in bacterial classification.

The sequencing of the 16S rRNA gene is commonly utilized to survey microbial communities, with specialized databases like Greengenes, RDP, and SILVA developed to support this method. It is recognized that most taxonomic annotations in these databases are predictions derived from sequences, rather than authoritative classifications based on studies of type strains or isolates (Edgar, 2018). Furthermore, it is acknowledged that databases like Greengenes and SILVA contain a considerable number of taxonomies that lack species-level resolution, which has limited the performance of classifiers (Hsieh et al., 2022).

The shortcomings of 16S rRNA sequencing extend beyond *Escherichia* and *Shigella*. Many bacterial genera, such as *Pseudomonas, Aeromonas*, and *Streptococcus*, exhibit similar problems where 16S rRNA-based classifications lack the discriminatory power to delineate species accurately (Jenkins et al., 2012; Janda and Abbott, 2007). The broader implications of this taxonomic debate highlight the need for standardized frameworks in bacterial classification. As 16S rRNA databases like Greengenes, SILVA, and RDP continue to evolve, taxonomic inconsistencies, particularly in closely related species like *Shigella* and *E. coli*, pose ongoing challenges. Therefore, a unified nomenclature system is essential for ensuring consistency and interoperability in bacterial taxonomy (Edgar, 2018).

Since querying the isolate’s 16S rRNA sequence and genome in recognised bacterial taxonomic databases did not yield a definitive or consistent taxonomic classification, Sourmash was used to cluster the isolate with additional genomes of bacterial species identified as close hits in the 16S rRNA NCBI BLAST. The isolate’s ambiguous clustering with both *Escherichia* and *Shigella* in Sourmash and phylogenetic analyses (Figures 2 and 3) highlights the limitations of relying solely on genomic approaches for distinguishing closely related species.

As several genomic approaches failed to yield a definitive and consistent taxonomic classification of the isolates, the biochemical and chromogenic differences among *E. coli, E. fergusonii*, and *Shigella* species on CCA were used to confirm the isolates’ identity. The isolate’s unique dark blue/purple appearance on CCA confirmed it as an *E. coli* strain, distinguishing it from *E. fergusonii* and *Shigella* species (Figure 6). This highlights the importance of simple laboratory biochemical and/or cultural procedures for identifying and differentiating *E. coli* from *Shigella* species, as well as from *E. fergusonii*.

*E. coli*, a coliform bacterium, ferments lactose and appears dark blue/purple on CCA. *E. fergusonii*, on the other hand, is also a coliform but does not ferment lactose, appearing pink on CCA due to its inability to hydrolyse X-glucuronide in the medium. *E. coli* and *E. fergusonii* are closely related bacteria, both capable of causing illnesses in humans, such as bacteraemia, urinary tract infections, and diarrhoea (Lindsey et al., 2017). *Shigella* species are non-coliform bacteria, do not ferment lactose, and appear colourless on CCA as they do not hydrolyse either Salmon-GAL or X-glucuronide in the medium. *E. fergusonii* has long been recognized as a close relative of bacteria in the *Escherichia coli*-*Shigella* group (Farmer et al., 1985), and the genetic similarities between *E. coli, E. fergusonii*, and *Shigella* species may present challenges for clinical diagnosis and effective treatment if phenotypic approaches, such as biochemical and cultural properties, are neglected. Therefore, integrating traditional microbiological approaches, including culturing and biochemical characterization methods, alongside MALDI-TOF mass spectrometry, to support the genomic identification of clinically significant bacteria is essential.

## CONCLUSION

This study describes the application of phenotypic and genomic approaches for the identification and antimicrobial characterization of an MDR bacterial strain, *E. coli* 266631E, isolated from the blood of a sepsis patient. The isolation of an MDR strain from a sepsis patient highlights the urgent need for diagnostic strategies that capture both genetic and phenotypic resistance, enabling timely and targeted antimicrobial therapy in life-threatening infections.

Initially identified as *E. coli* through MALDI-TOF analysis, the isolate exhibited an alarming MDR profile after phenotypic antimicrobial screening (disk diffusion testing and broth microdilution MIC determination), prompting genomic studies to confirm its taxonomic identity and characterize its ARGs. Genomic analysis revealed multiple ARGs encoding resistance to a broad range of clinically important antibiotics across different classes, including macrolides, fluoroquinolones, aminoglycosides, carbapenems, and cephalosporins. However, attempts to resolve the isolate’s taxonomy by querying its 16S rRNA and genome sequences in recognised bacterial taxonomic databases, such as NCBI, Greengenes2, and SILVA, were inconclusive. The conserved nature of the 16S rRNA gene and the close phylogenetic relationship between *Shigella* and *Escherichia* hindered definitive species-level classification.

Although *E. coli* is extensively studied, its taxonomic classification remains challenging due to high genetic similarity with species like *Shigella* and *E. fergusonii*, particularly in conserved genomic regions—despite their phenotypic differences and variations in pathogenic traits. The isolate’s ambiguous clustering with both *Escherichia* and *Shigella* in Sourmash and phylogenetic analyses highlights the limitations of relying solely on genomic methods for distinguishing closely related taxa. Ultimately, the species identity of *E. coli* 266631E was confirmed through a simple phenotypic assay using CCA, emphasizing the continued relevance of traditional microbiological techniques alongside modern genomic tools.

This study underscores the importance of integrating phenotypic methods with genomic analyses for accurate identification of clinically significant bacteria. Such integration will enhance effective surveillance of key MDR variants in healthcare settings, enabling precise diagnosis and effective treatment of life-threatening infections such as sepsis.

