## Supplementary Material for "Whole-Genome Sequencing, Annotation and Phenotypic Taxonomic Confirmation of a Multidrug-Resistant *Escherichia coli* Strain Isolated from the Blood of a Sepsis Patient"

**TABLE OF CONTENTS**

| Content | | Page |
| --- | --- | --- |
| Figure S1: | Result of MALDI-TOF analysis | 3 |
| Figure S2: | Screenshot of AMRFinderPlus output showing AMR genes in contig 1 (chromosome) and contig 3 (IncF-type plasmid) showing predicted AMR genes at 100% identity | 4 |
| Figure S3: | Screenshot of starAMR output showing AMR genes in contig 1 (chromosome) and contig 3 (IncF-type plasmid) showing predicted AMR genes at ≥99% identity | 5 |
| Figure S4: | Screenshot of RGI output of contig 1 (chromosome) showing predicted AMR genes at ≥99% identity | 6 |
| Figure S5: | Screenshot of RGI output of contig 3 (IncF-type plasmid) showing predicted AMR genes at ≥99% identity | 9 |
| Figure S6: | NCBI BLAST Search of bacterial 16S rRNA sequence on NCBI Standard Databases [Core nucleotide database option (core nt) option] | 10 |
| Figure S7: | NCBI BLAST Search of bacterial 16S rRNA sequence on NCBI’s rRNA/ITS databases [16S ribosomal RNA sequences (Bacterial and Archaea) option] | 11 |
| Figure S8: | Snapshot of Greengenes2 BLAST result output | 12 |
| Figure S9: | Snapshot of SILVA BLAST result output | 13 |
| Figure S10: | Sourmash of 1,595 genomes showing the average nucleotide identity (ANI) | 14 |
| Figure S11: | Sourmash of 1,595 genomes showing the Jaccard index | 15 |
| Figure S12: | (A) Phylogeny of the isolate using FastTree (B) Reduced representation of 1,595 isolates | 16 |

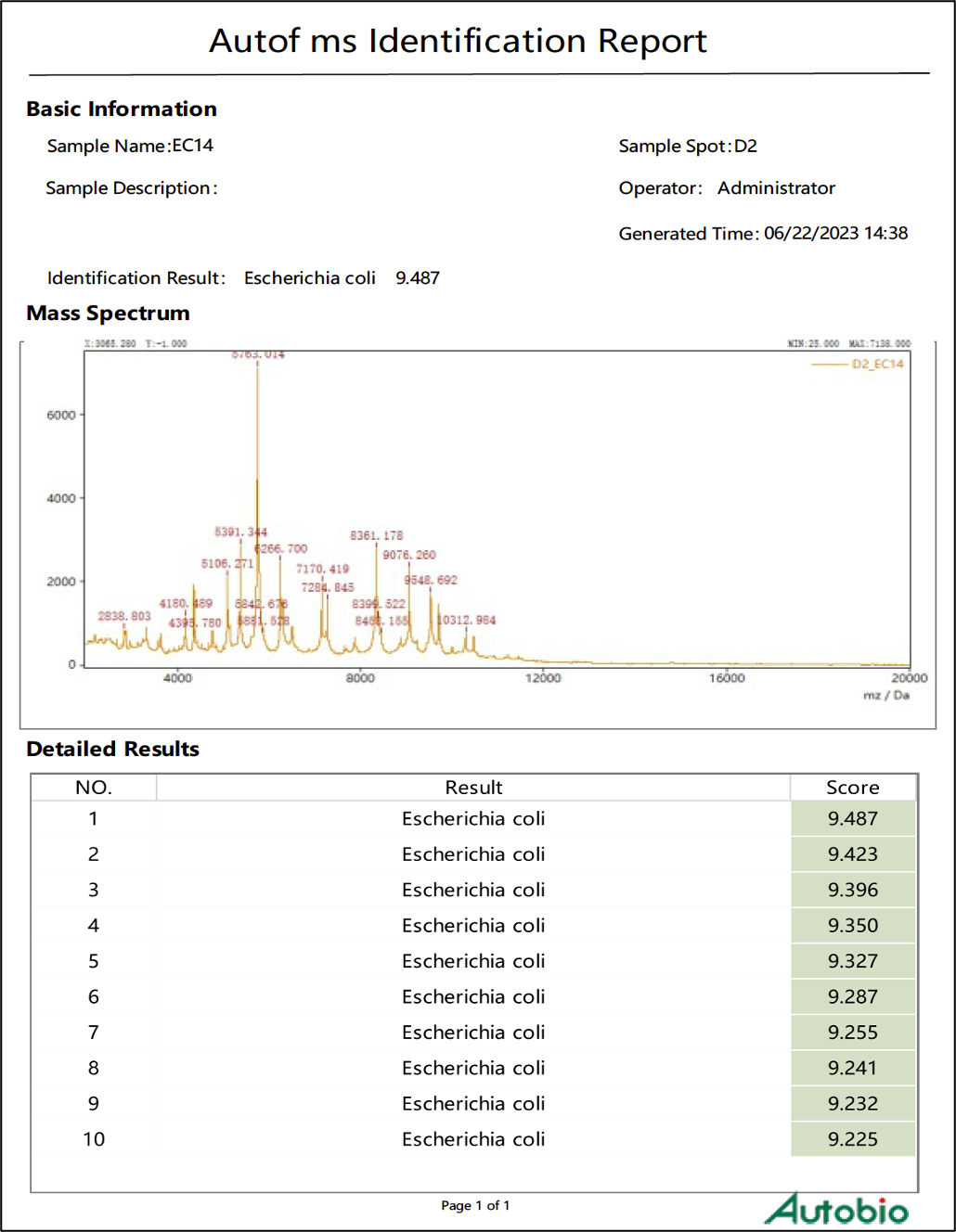

**Figure S1:** Result of MALDI-TOF analysis

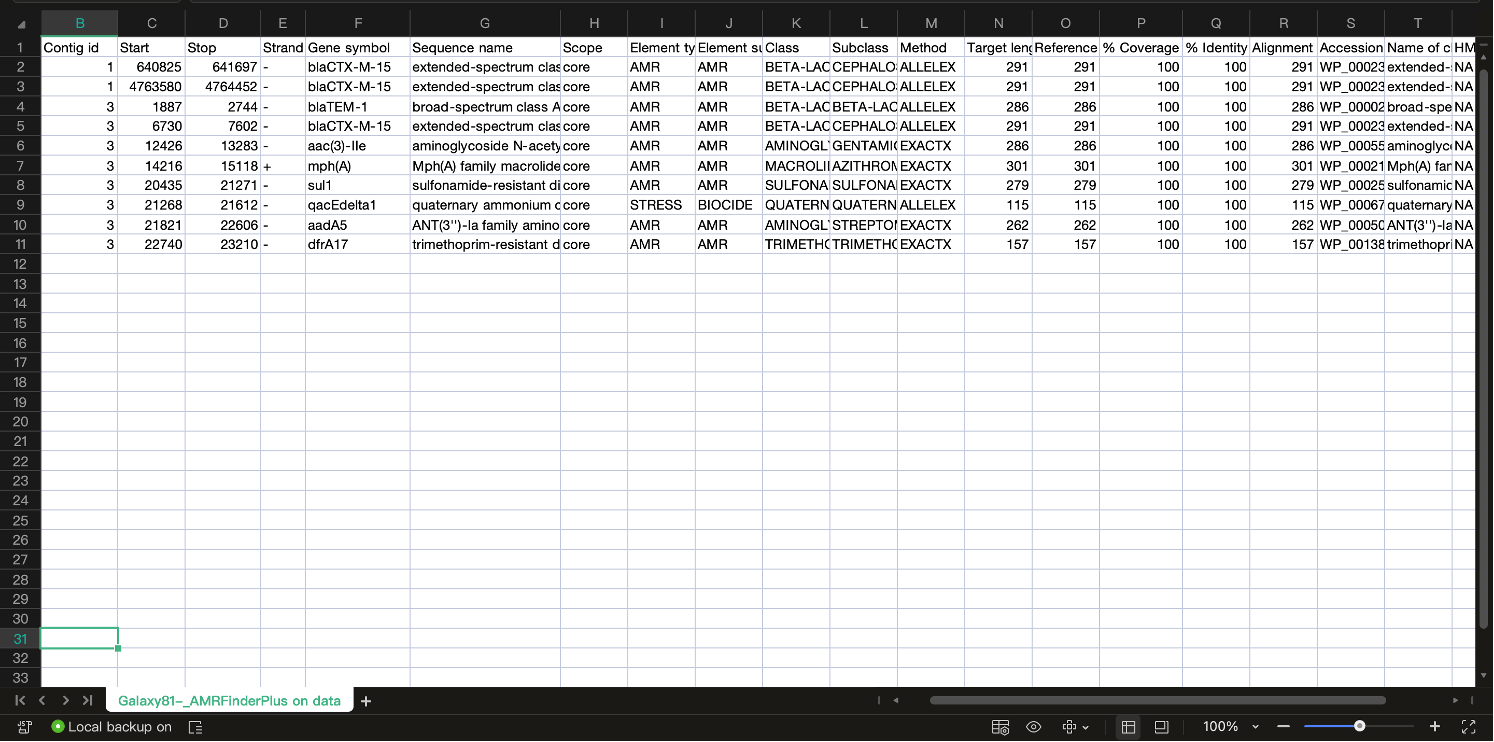

**Figure S2:** Screenshot of AMRFinderPlus output showing AMR genes in contig 1 (chromosome) and contig 3 (IncF-type plasmid) showing predicted AMR genes at 100% identity

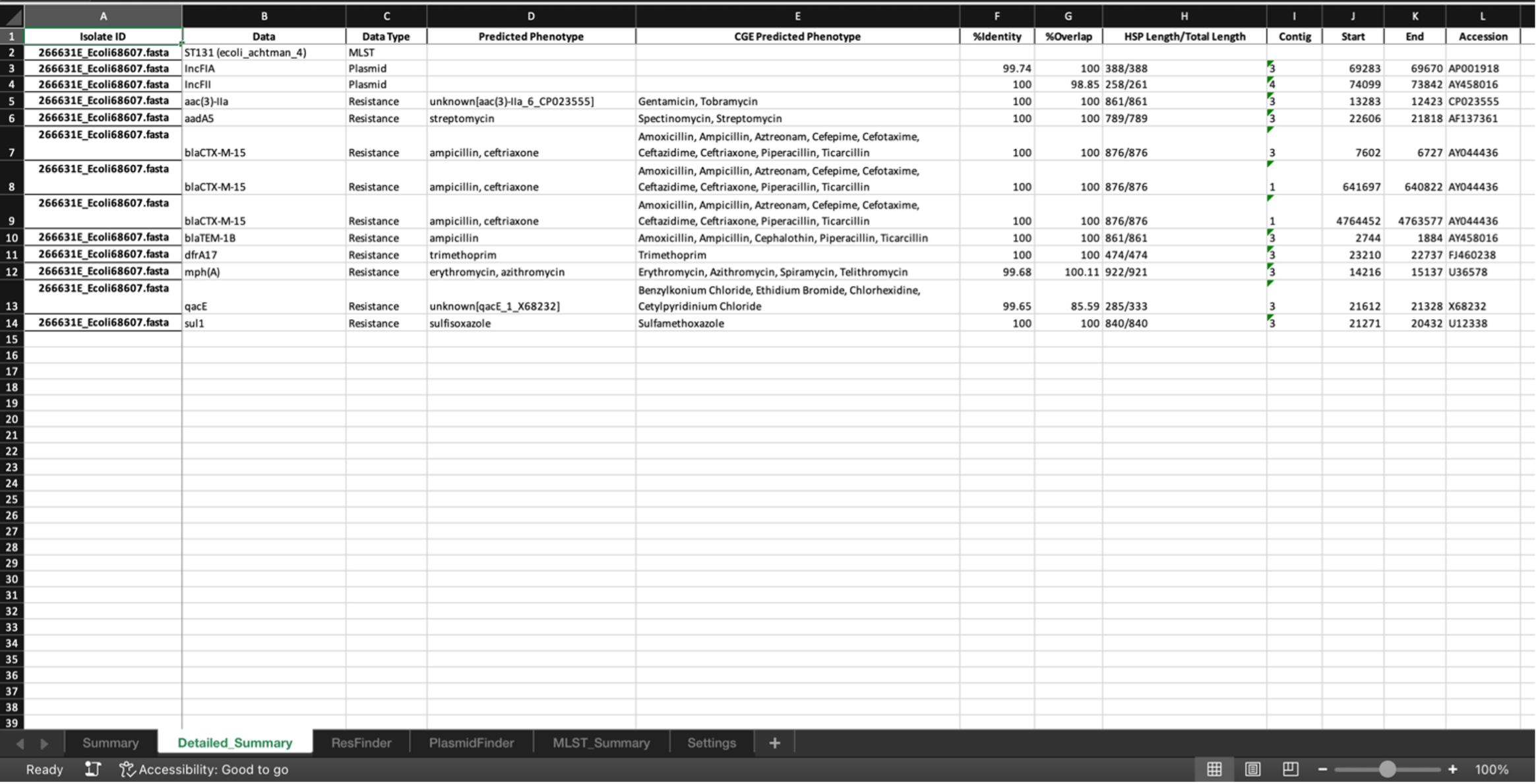

**Figure S3:** Screenshot of starAMR output showing AMR genes in contig 1 (chromosome) and contig 3 (IncF-type plasmid) showing predicted AMR genes at ≥99% identity

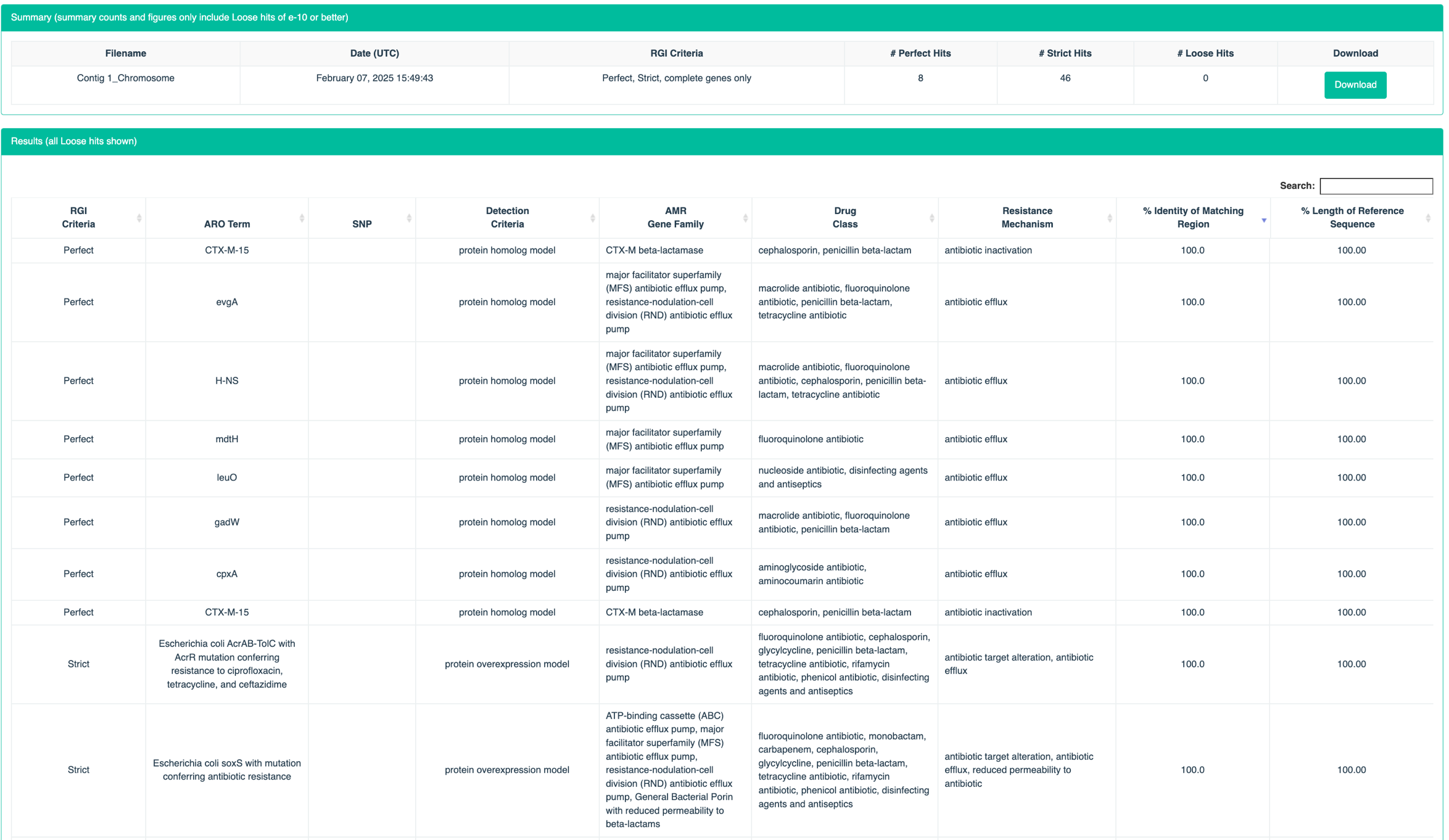

**Figure S4:** Screenshot of RGI output of contig 1 (chromosome) showing predicted AMR genes at ≥99% identity

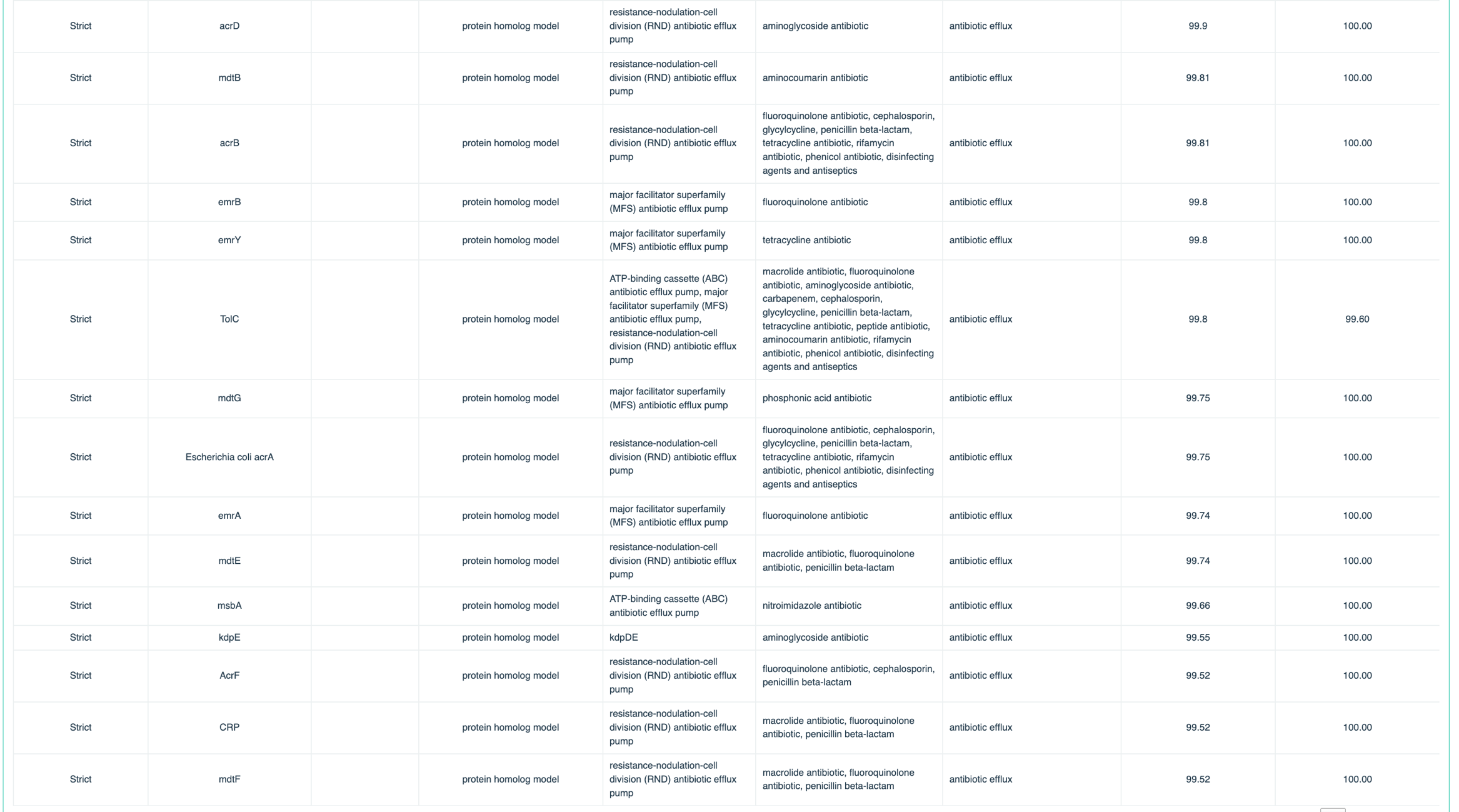

**Figure S4:** Screenshot of RGI output of contig 1 (chromosome) showing predicted AMR genes at ≥99% identity…(continued_2)

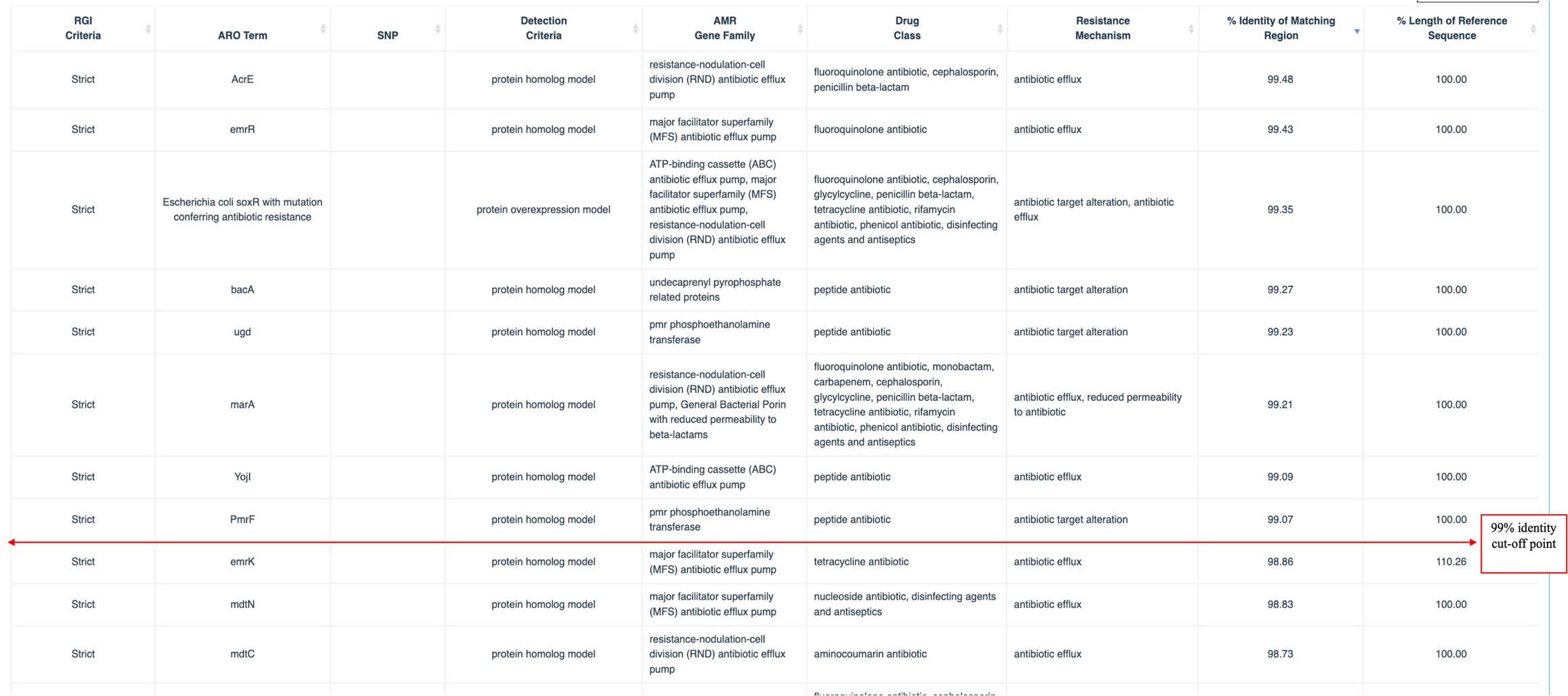

**Figure S4:** Screenshot of RGI output of contig 1 (chromosome) showing predicted AMR genes at ≥99% identity…(continued_3)

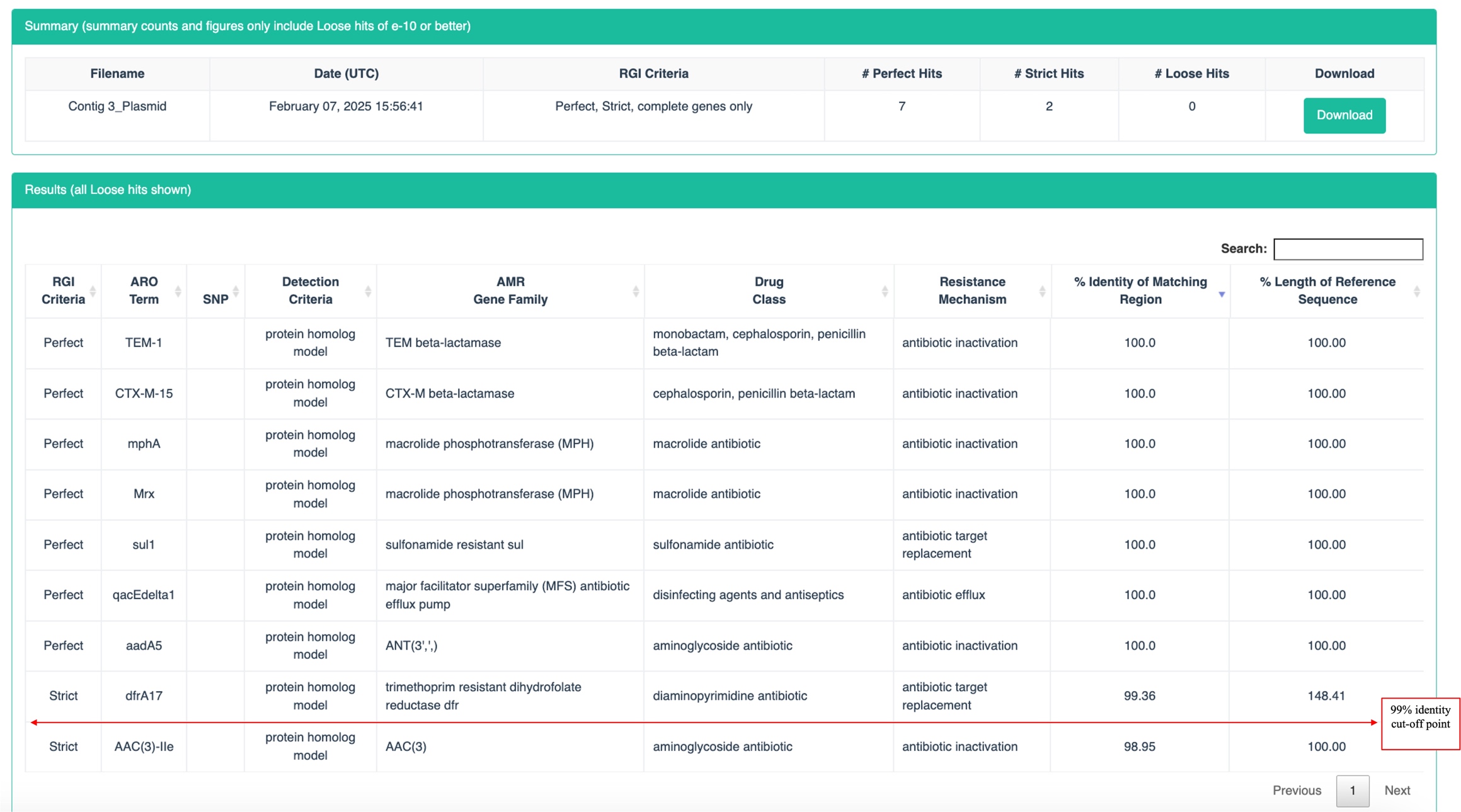

**Figure S5:** Screenshot of RGI output of contig 3 (IncF-type plasmid) showing predicted AMR genes at ≥99% identity

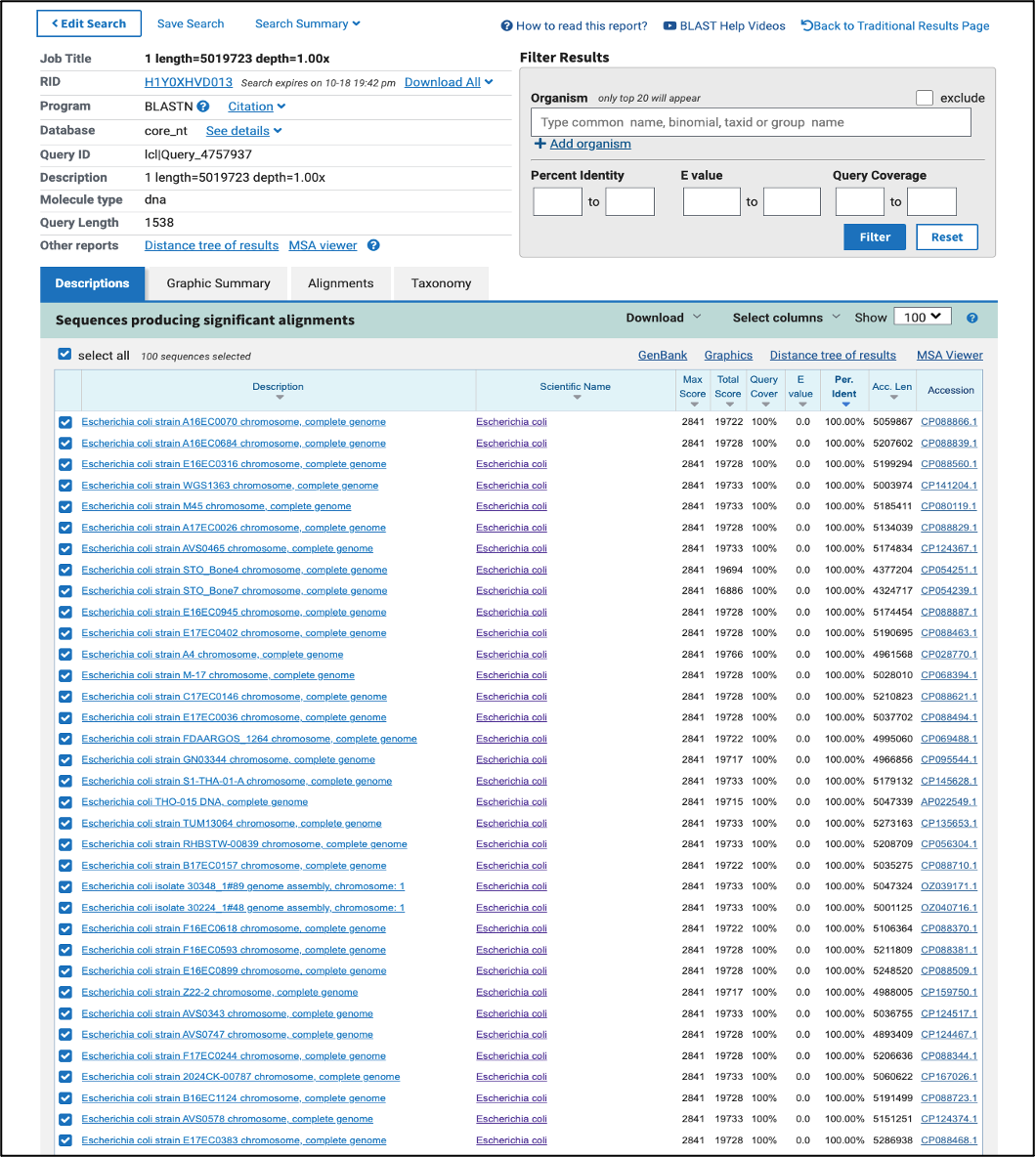

**Figure S6:** NCBI BLAST Search of bacterial 16S rRNA sequence on NCBI Standard Databases [Core nucleotide database option (core nt) option]

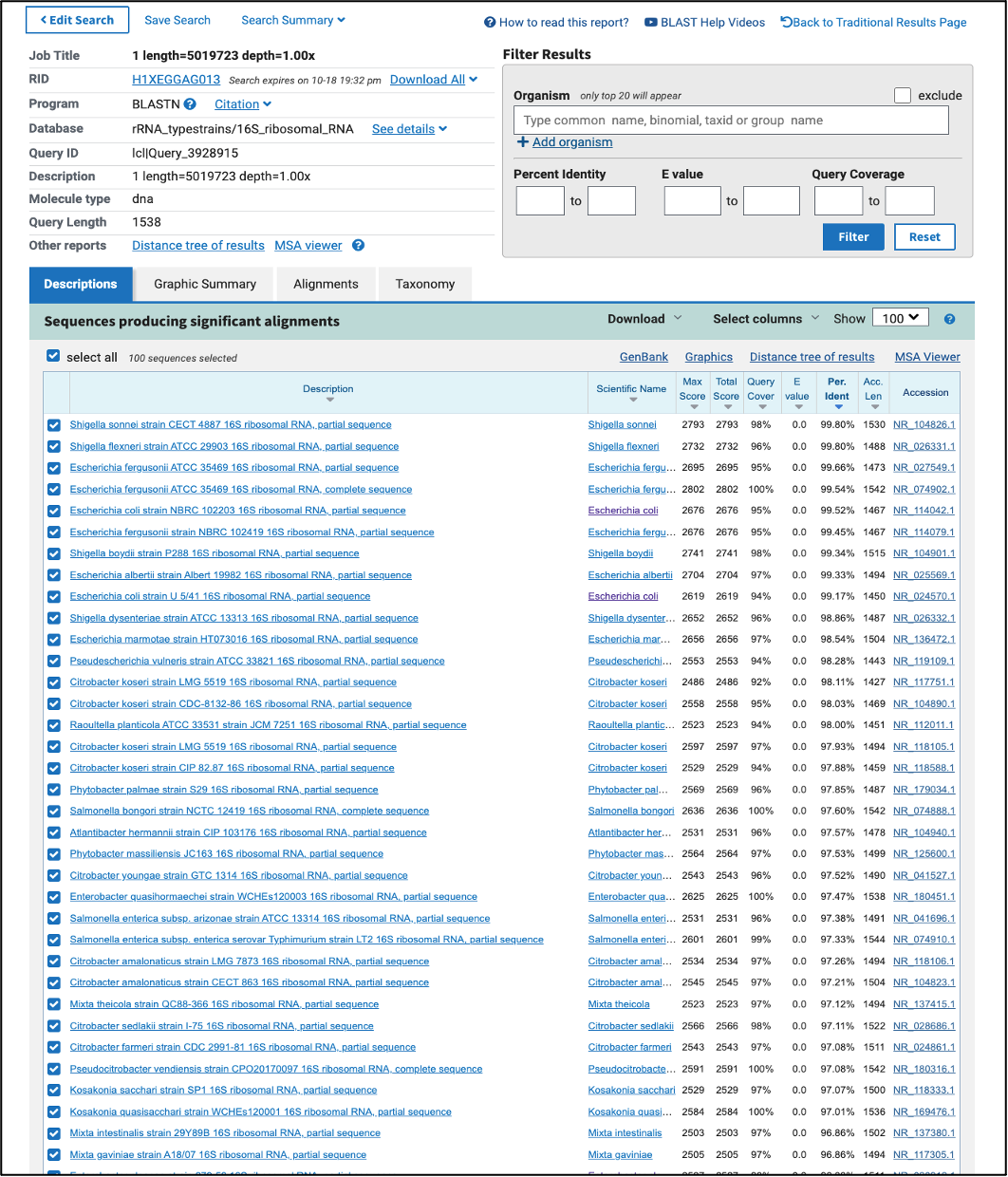

**Figure S7:** NCBI BLAST Search of bacterial 16S rRNA sequence on NCBI’s rRNA/ITS databases [16S ribosomal RNA sequences (Bacterial and Archaea) option]

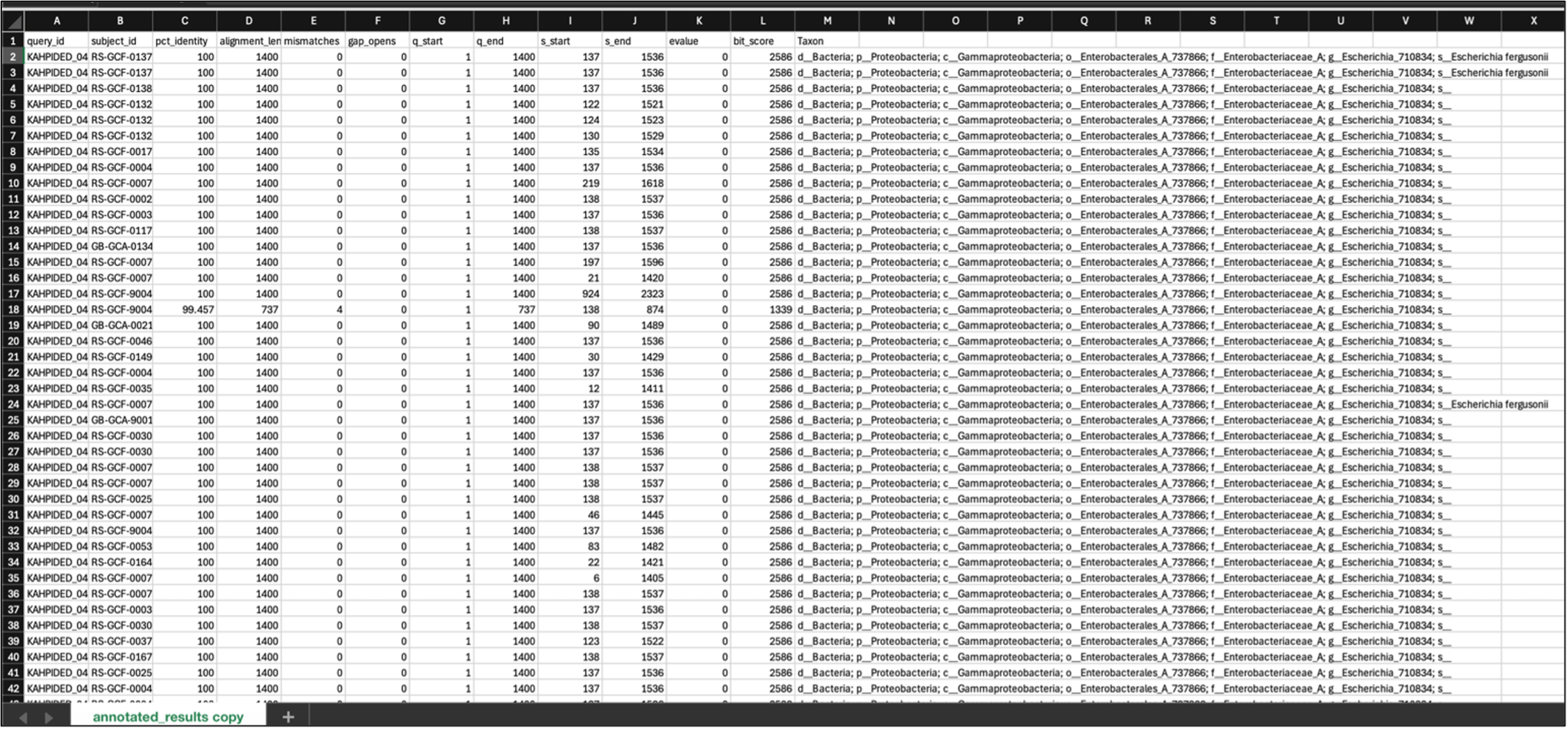

**Figure S8:** Snapshot of Greengenes2 BLAST result output

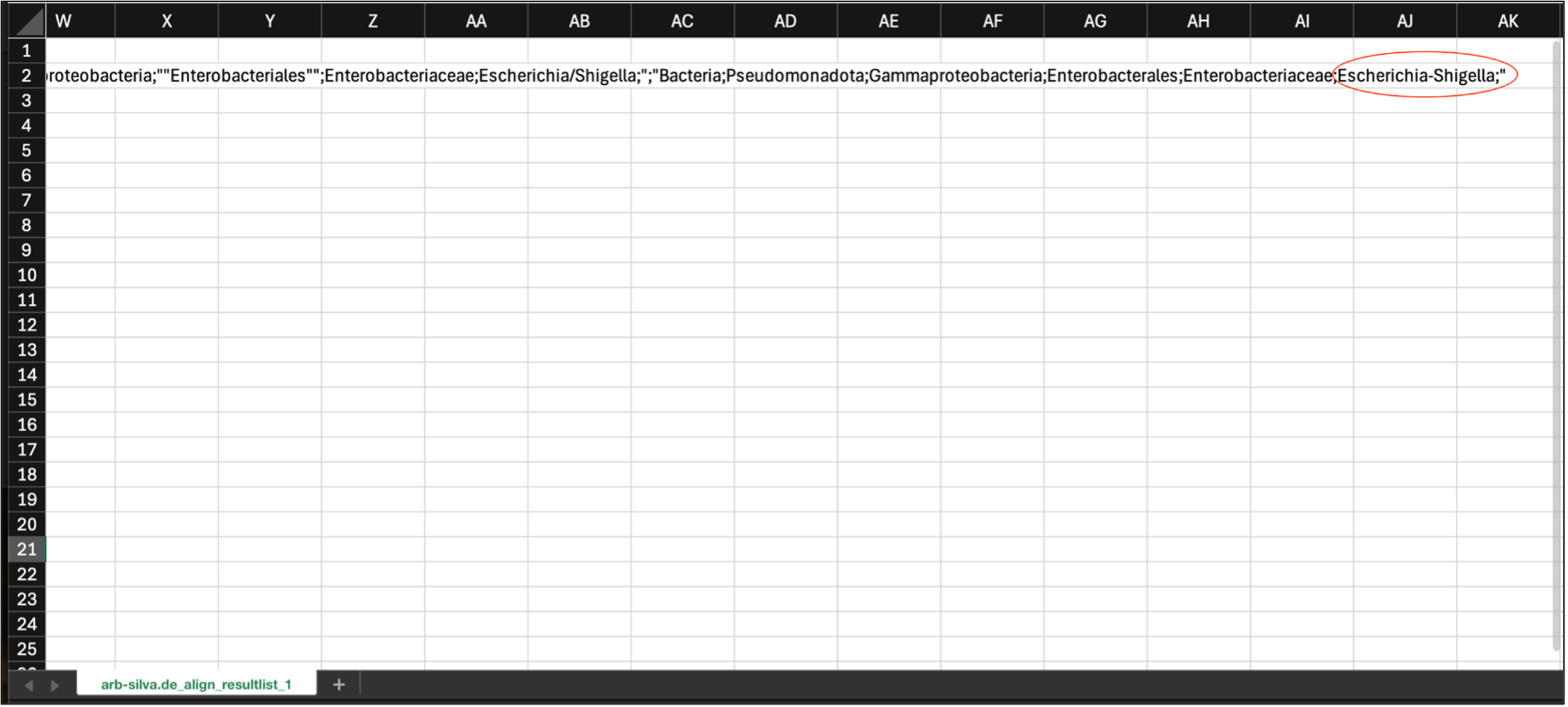

**Figure S9:** Snapshot of SILVA BLAST result output

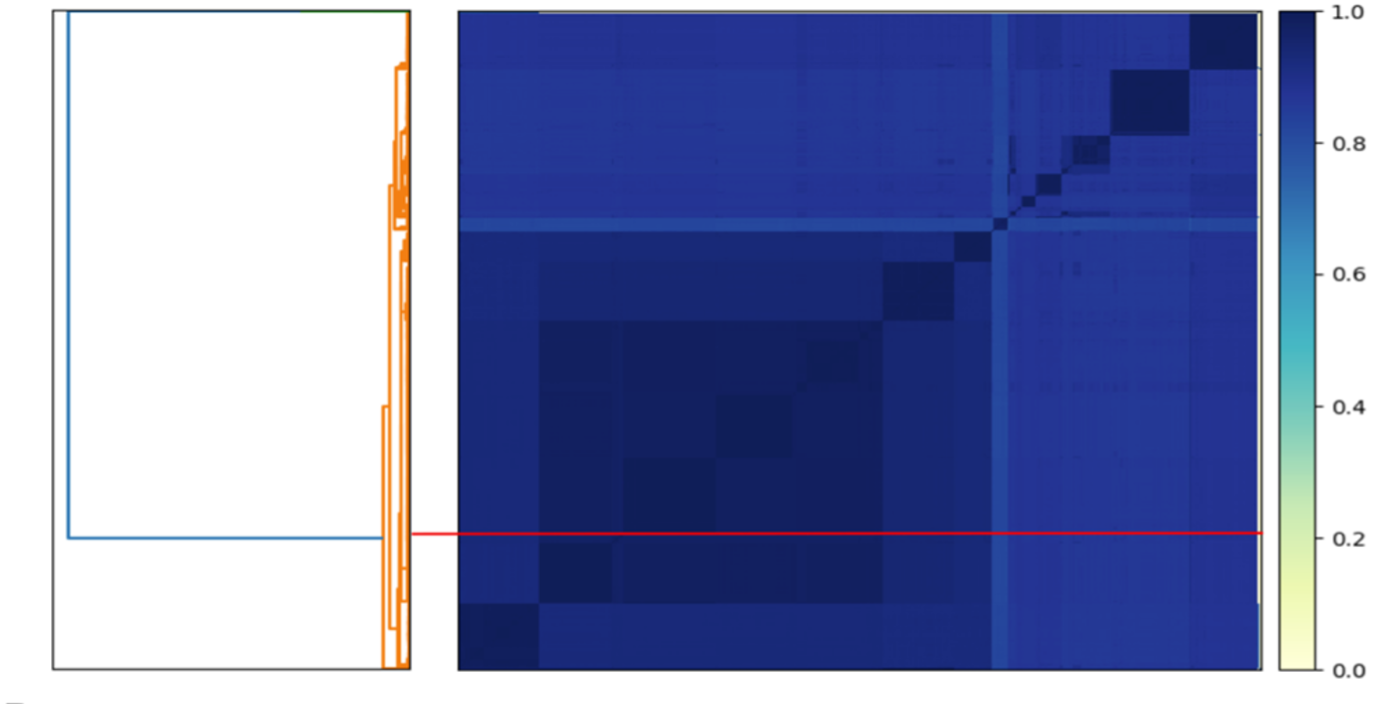

**Figure S10:** Sourmash of 1,595 genomes showing the average nucleotide identity (ANI). The red line on both plots indicates the placement of the isolate

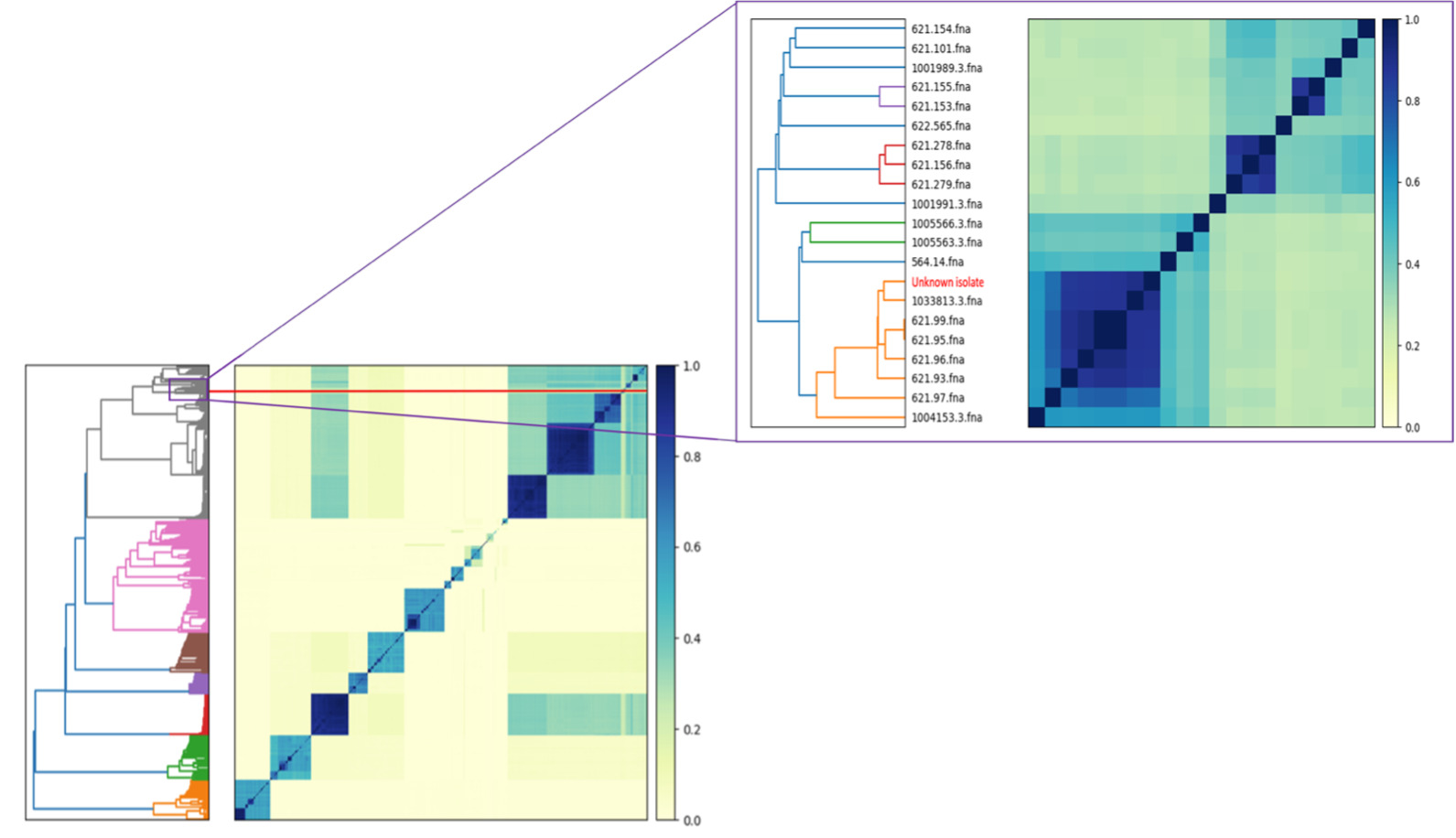

**Figure S11:** Sourmash of 1,595 genomes showing the Jaccard index. The darker the colour, the more genetically similar the isolates are. The red line in the plot indicates the placement of the isolate (labelled as ''unknown isolate''), which is clustered among multiple *Shigella* and *Escherichia* spp. Note: Genome IDs beginning with 621 belong to the genus *Shigella*, while the others belong to *Escherichia*.

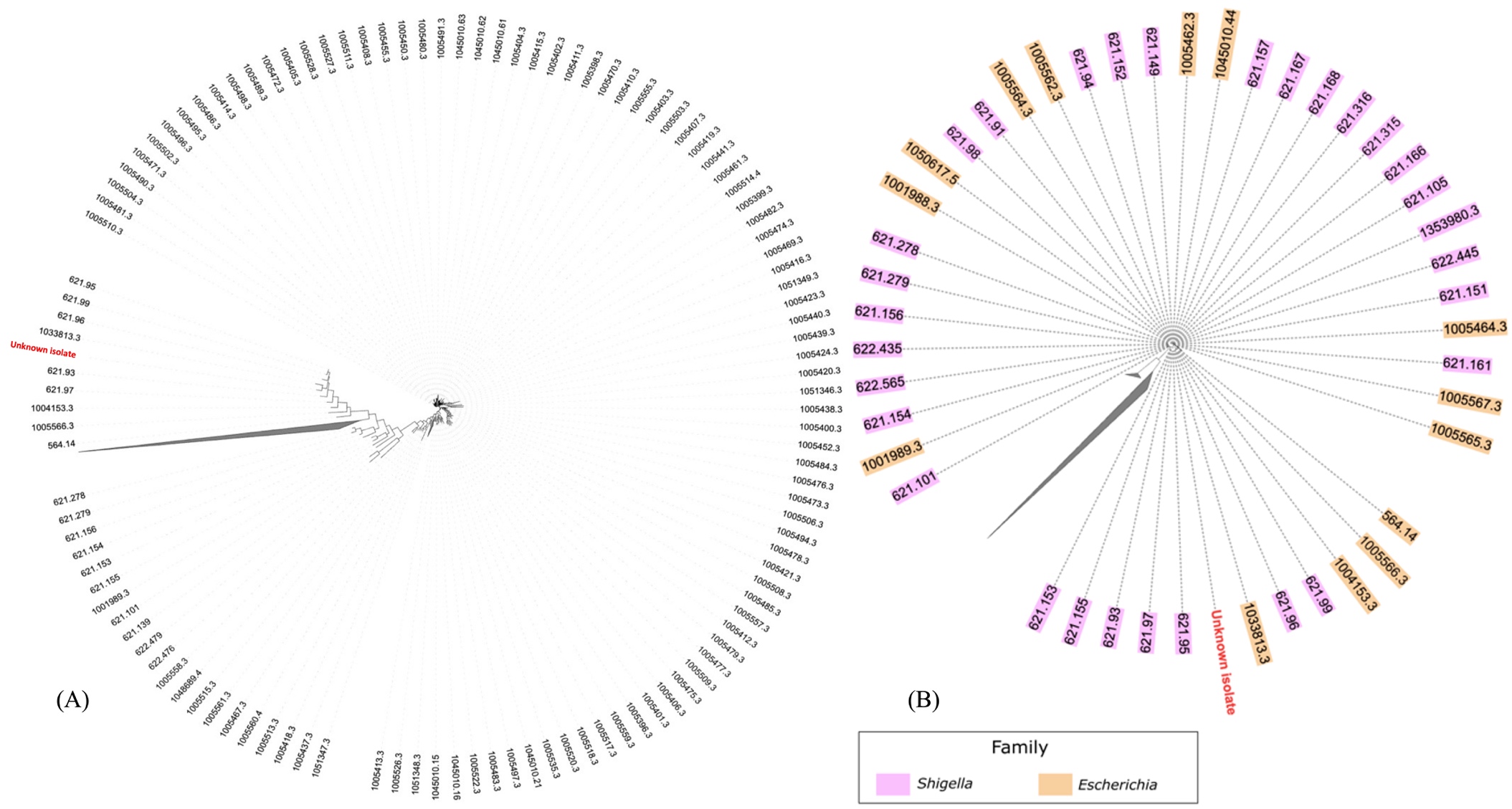

**Figure S12:** (A) Phylogeny of the isolate using FastTree (B) Reduced representation of 1,595 isolates (clades have been removed to be able to visualise the isolate and surrounding genomes).
